# Impact of SARS-CoV-2 ORF6 and its variant polymorphisms on host responses and viral pathogenesis

**DOI:** 10.1101/2022.10.18.512708

**Authors:** Thomas Kehrer, Anastasija Cupic, Chengjin Ye, Soner Yildiz, Mehdi Bouhhadou, Nicholas A Crossland, Erika Barrall, Phillip Cohen, Anna Tseng, Tolga Çağatay, Raveen Rathnasinghe, Daniel Flores, Sonia Jangra, Fahmida Alam, Nacho Mena, Sadaf Aslam, Anjali Saqi, Arturo Marin, Magdalena Rutkowska, Manisha R. Ummadi, Giuseppe Pisanelli, R. Blake Richardson, Ethan C. Veit, Jacqueline M. Fabius, Margaret Soucheray, Benjamin J. Polacco, Matthew J. Evans, Danielle L. Swaney, Ana S. Gonzalez-Reiche, Emilia M. Sordillo, Harm van Bakel, Viviana Simon, Lorena Zuliani-Alvarez, Beatriz M. A. Fontoura, Brad R. Rosenberg, Nevan J. Krogan, Luis Martinez-Sobrido, Adolfo García-Sastre, Lisa Miorin

## Abstract

We and others have previously shown that the SARS-CoV-2 accessory protein ORF6 is a powerful antagonist of the interferon (IFN) signaling pathway by directly interacting with Nup98-Rae1 at the nuclear pore complex (NPC) and disrupting bidirectional nucleo-cytoplasmic trafficking. In this study, we further assessed the role of ORF6 during infection using recombinant SARS-CoV-2 viruses carrying either a deletion or a well characterized M58R loss-of-function mutation in ORF6. We show that ORF6 plays a key role in the antagonism of IFN signaling and in viral pathogenesis by interfering with karyopherin(importin)-mediated nuclear import during SARS-CoV-2 infection both *in vitro*, and in the Syrian golden hamster model *in vivo*. In addition, we found that ORF6-Nup98 interaction also contributes to inhibition of cellular mRNA export during SARS-CoV-2 infection. As a result, ORF6 expression significantly remodels the host cell proteome upon infection. Importantly, we also unravel a previously unrecognized function of ORF6 in the modulation of viral protein expression, which is independent of its function at the nuclear pore. Lastly, we characterized the ORF6 D61L mutation that recently emerged in Omicron BA.2 and BA.4 and demonstrated that it is able to disrupt ORF6 protein functions at the NPC and to impair SARS-CoV-2 innate immune evasion strategies. Importantly, the now more abundant Omicron BA.5 lacks this loss-of-function polymorphism in ORF6. Altogether, our findings not only further highlight the key role of ORF6 in the antagonism of the antiviral innate immune response, but also emphasize the importance of studying the role of non-spike mutations to better understand the mechanisms governing differential pathogenicity and immune evasion strategies of SARS-CoV-2 and its evolving variants.

**ONE SENTENCE SUMMARY:** SARS-CoV-2 ORF6 subverts bidirectional nucleo-cytoplasmic trafficking to inhibit host gene expression and contribute to viral pathogenesis.

## MAIN TEXT

Despite the rapid development of vaccines and antiviral treatments, the coronavirus disease 2019 (COVID-19) pandemic, caused by severe respiratory syndrome coronavirus 2 (SARS-CoV-2), still remains a major global health concern (https://covid19.who.int). The clinical presentations of COVID-19 involve a broad range of symptoms, from asymptomatic infections to severe disease, normally characterized by excessive induction of proinflammatory cytokines, with an overall fatality rate near 1% (*1, 2*). While the determinants for disease outcome are not completely understood, numerous studies have suggested that the inability to mount a timely and effective antiviral IFN response promotes viral persistence and tissue damage, contributing to SARS-CoV-2 virulence and COVID-19 severity (*1, 3, 4*). In this regard, inborn errors of immunity affecting the TLR3 or IFN pathway (ref), and the presence of neutralizing autoantibodies against type I IFN (*5, 6*), have been identified in a subset of severe COVID-19 patients. Furthermore, several viral proteins have been described to inhibit or suppress innate immune activation at different levels (*7–9*), highlighting the importance of type I IFN in the defense against SARS-CoV-2 infection. Among these proteins, the non-structural protein NSP1 has been shown to inhibit antiviral gene expression by inhibiting translation (*10*), blocking nuclear export of cellular transcripts (*11, 12*), and inducing host mRNA cleavage (*13*). The accessory protein ORF9B antagonizes IFN induction by interacting with TOM70 and inhibiting mitochondrial recruitment of TBK1 and IRF3 (*14*). In addition, SARS-CoV-2 ORF6 was found to directly interact with the Nup98-Rae1 complex to disrupt karyopherin-mediated nuclear import of STAT1 and STAT2 (*8*), and to contribute to the inhibition of mRNA export that we and others have observed during infection (*15–17*).

As the virus evolved since its initial introduction into humans, new SARS-CoV-2 variants have emerged with major genomic changes that confer resistance to neutralizing antibodies and exhibit increase transmissibly and virulence (*18, 19*). Remarkably, we have previously shown that such variants of concern (VOCs), in addition to gain spike mutations that mediate antibody escape and alter virus entry into human cells, also evolved non-spike mutations that result in increased expression of key viral innate immune antagonists such as ORF9B and ORF6, and enhanced innate immune suppression (*20*). In this study, we closely dissect the impact of ORF6 and its recently emerged variant polymorphisms on the host response to SARS-CoV-2 infection to gain more detailed insights into the mechanisms employed by SARS-CoV-2 to escape innate antiviral responses and drive COVID-19 pathogenesis.

## RESULTS

### ORF6 expression is essential for inhibition of STAT nuclear import during infection with SARS-CoV-2

We and others have previously shown that the SARS-CoV-2 accessory protein open reading frame 6 (ORF6) directly interacts with Nup98-Rae1 at the nuclear pore complex (NPC) to disrupt STAT nuclear translocation and antagonize IFN signaling (*8, 9*). In this study, we employed our previously described recombinant SARS-CoV-2 virus system (*21–23*) to further assess the role of ORF6 in the modulation of the innate immune response in the context of infection. As shown in Fig. 1A, in addition to a recombinant SARS-CoV-2 wildtype virus (rSARS-CoV-2 WT), we generated a mutant virus carrying a deletion of the ORF6 coding sequence (rSARS-CoV-2 ΔORF6), as well as a virus with the ORF6^M58R^ mutation (rSARS-CoV-2 ORF6^M58R^) previously shown to abolish binding to the Nup98-Rae1 complex (*8*). The presence of the ORF6 deletion and ORF6^M58R^ mutation were validated by genome sequencing of the viral stocks (Fig. S1). Next, to evaluate differences in viral growth, we monitored the replication kinetics of the different recombinant viruses in both Vero E6 and A549-ACE2 cells. Interestingly, while infection of Vero E6 cells did not reveal significant differences in viral titers at any of the time points analyzed, we found that both the ORF6-deficient and the ORF6^M58R^ viruses replicated to lower titers than the wildtype virus in IFN-competent A549-ACE2 cells (Fig. 1B-C). As we previously showed that ORF6 antagonizes IFN signaling downstream of STAT phosphorylation (*1*), we then assessed the ability of the ORF6 mutant viruses to inhibit STAT phosphorylation and nuclear translocation. As expected, upon treatment of control or infected Vero E6 cells with recombinant IFN, we observed no differences in the levels of total or phosphorylated STAT1 and STAT2 across conditions (Fig. 1D). However, immunofluorescence microscopy revealed that IFN-dependent STAT2 nuclear translocation was effectively rescued in cells infected with both the ORF6-deficient and the ORF6^M58R^ viruses (Fig. 1F). Importantly, these results were also confirmed in A549-ACE2 cells that can endogenously trigger IFN induction and subsequent STAT phosphorylation and nuclear translocation in response to SARS-CoV-2 infection (Fig. 1E and G). Of note, A549-ACE2 cells showed similar levels of STAT1 and STAT2 phosphorylation upon infection with the wildtype or ORF6 mutant viruses, suggesting that while ORF6 expression plays a major role in the antagonism of IFN signaling, its role in the inhibition of IFN induction during infection might be redundant.

**Figure 1.**
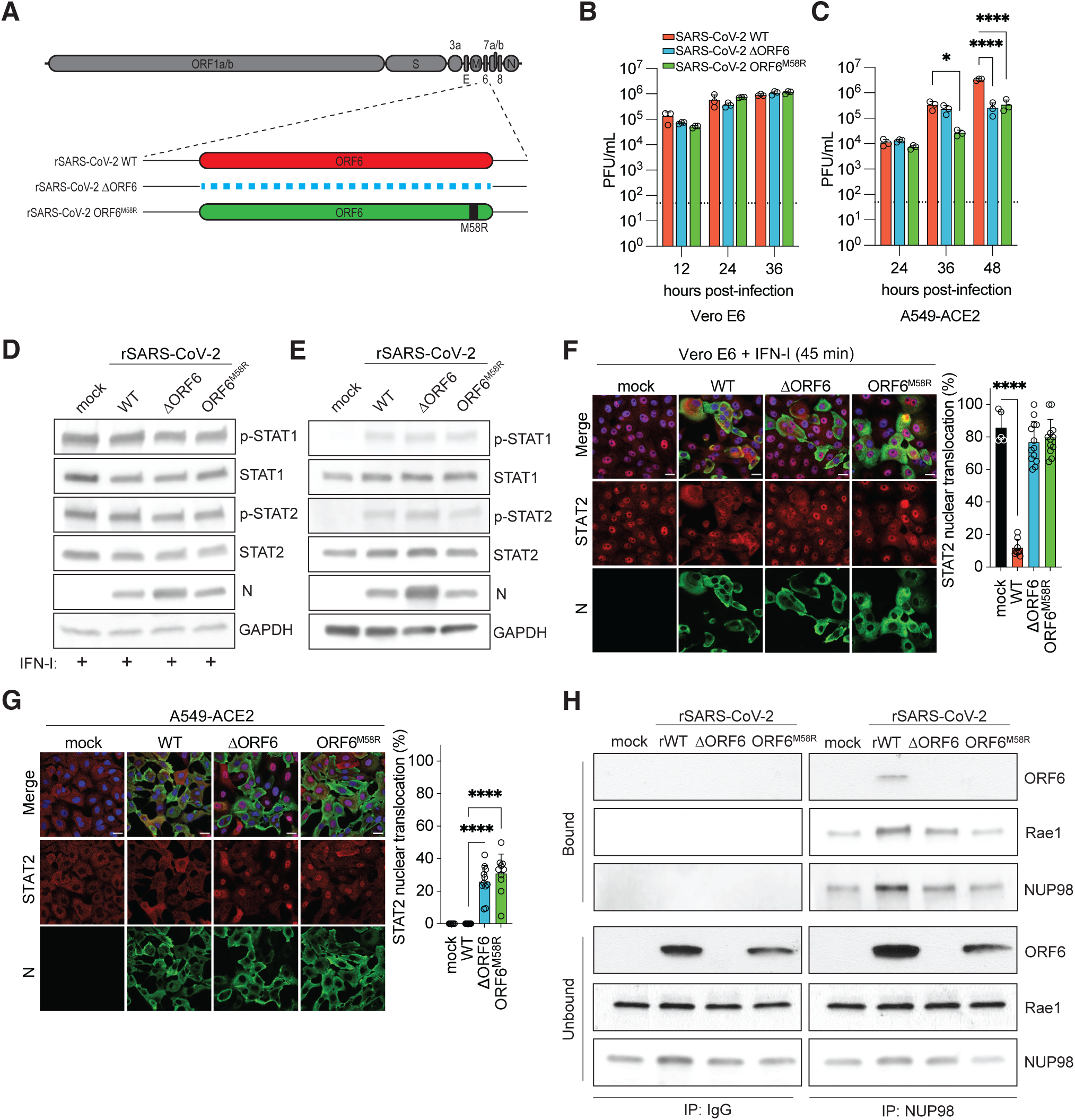
ORF6 is essential for inhibition of STAT nuclear import and optimal SARS-CoV-2 replication in IFN competent cells. (A) Schematic illustration of the genome organization of rSARS-CoV-2 WT, ΔORF6, and ORF6^M58R^ viruses used in our studies. (B) Growth curve of rSARS-CoV-2 WT, ΔORF6, and ORF6^M58R^ in Vero E6 cells infected at MOI 0.1 for 12, 24, and 36 hours. Titers were quantified by plaque assay (n=3). (C) Growth curve of rSARS-CoV-2 WT, ΔORF6, and ORF6^M58R^ in A549-ACE2 cells infected at MOI 0.1 for 12, 24, and 36hours. Titers were quantified by plaque assay (n=3). (D) Vero E6 cells were either mock-infected or infected with the indicated recombinant viruses at MOI 0.5 for 24 hours. Cells were treated with 1000 U of IFN universal for 45 min prior to lysis and used for western blot analysis. Expression and phosphorylation status of the indicated proteins was determined by Western Blot using GAPDH as a loading control. (E) A549-ACE2 cells were either mock-infected or infected with the indicated recombinant viruses at MOI 0.5 for 24 hours. Expression and phosphorylation status of the of the indicated proteins was determined by Western Blot using GAPDH as a loading control. (F) Confocal microscopy images of Vero E6 cells infected with the indicated viruses at MOI 0.5 for 24 hours. Cells were treated with 1000 U of IFN universal for 45 min prior to fixation. The subcellular localization of STAT2 was analyzed by confocal microscopy. Nuclei were stained with DAPI. STAT2 nuclear translocation in infected and bystander cells was quantified from ≥150 cells per condition from two biological replicates and compared to translocation in mock-infected cells. Data are shown as average ± SD. (G) Confocal microscopy images of A549-ACE2 cells infected with the indicated viruses at MOI 0.5 for 24 hours prior to fixation. The subcellular localization of STAT2 was analyzed by confocal microscopy. Nuclei were stained with DAPI. STAT2 nuclear translocation in infected and bystander cells was quantified from ≥150 cells per condition from two biological replicates and compared to translocation in mock-infected cells. Data are shown as average ± SD. (H) A549-ACE2 cells were either mock-infected or infected with the indicated viruses for 24 hours and the subjected to immunoprecipitation of endogenous Nup98 followed by western blot analysis to detect the indicated proteins. (Scale bar = 20 µm). Data in B and C were analyzed by two-way ANOVA using Dunnett’s multiple comparison test. Data in F and G were analyzed by ordinary one-way ANOVA using Turkey’s multiple comparison test. P < 0.05 = *, P < 0.0001 = ****. Graphs were generated with PRISM (version 9).

Finally, to confirm the interaction of ORF6 with the Nup98-Rae1 complex in the context of infection, we immunoprecipitated endogenous Nup98 in A549-ACE2 cells that were either mock infected or infected with the three different recombinant viruses. Strikingly, in agreement with our earlier findings, both ORF6 and Rae1 co-immunoprecipitated with Nup98 in cells infected with rSARS-CoV-2 WT, while only Rae1 was efficiently pulled-down by Nup98 in cells infected with both the ORF6-deficient and the ORF6^M58R^ viruses, as expected (Fig. 1H). All together, these results indicate that ORF6 binds to the Nup98-Rae1 complex during SARS-CoV-2 infection and that such virus-host interaction plays a major role in the antagonism of the IFN signaling pathway by disrupting STAT nuclear translocation.

### SARS-CoV-2 ORF6 selectively blocks nuclear import of transcription factors

Next, to further investigate the role of ORF6 in the subversion of other important signaling pathways involved in the host antiviral response, we closely looked at its ability to inhibit IRF3 and NFkB nuclear translocation. In agreement with previous findings (*8, 9*), we show that ectopic expression of ORF6, but not of ORF6^M58R^ that loses nuclear pore binding ability, was able to significantly block RIG-I-2CARD-mediated IRF3-GFP nuclear translocation (Fig. 2A) as well as IRF3-dependent gene expression in HEK293T cells as assessed by a reporter-based assay (Fig. 2C). However, p65 nuclear translocation and NFkB reporter activation upon TNFa treatment were not affected by overexpression of increasing amounts of ORF6 or ORF6^M58R^ (Fig. 2B and D). For these experiments, expression of Hepatitis C virus (HCV) NS3/4A and TRIM9 were used as positive control for inhibition of gene expression upstream of the IRF3- and NF-kB-responsive promoters, respectively (*24, 25*).To address the relevance of these findings in the context of infection, we then infected A549-ACE2 cells with either wildtype or ORF6-deficient SARS-CoV-2 at MOI 1 and quantified both IRF3 and NFkB nuclear translocation by immunofluorescence analysis. As expected, we found that p65 efficiently translocated into the nucleus of cells infected with both recombinant viruses (Fig. 2F). However, we did not find significant differences in IRF3 nuclear translocation at any of the time points analyzed (Fig. 2G, and data not shown). Importantly, these data were also consistent with the similar levels of IRF3 and NFkB phosphorylation detected by Western blot analysis of cells infected with the wildtype and ORF6-deficient viruses (Fig. 2E). This suggests that while ORF6 has the potential to block IRF3 nuclear translocation by interfering with karyopherin-mediated nuclear import, its function in the inhibition of IFN induction during infection is likely redundant. Presumably, this is due to the expression of other viral antagonists that are acting more upstream in the pathway, and contribute to the poor and delayed IRF3 activation by SARS-CoV-2 that we and others have observed.

**Figure 2.**
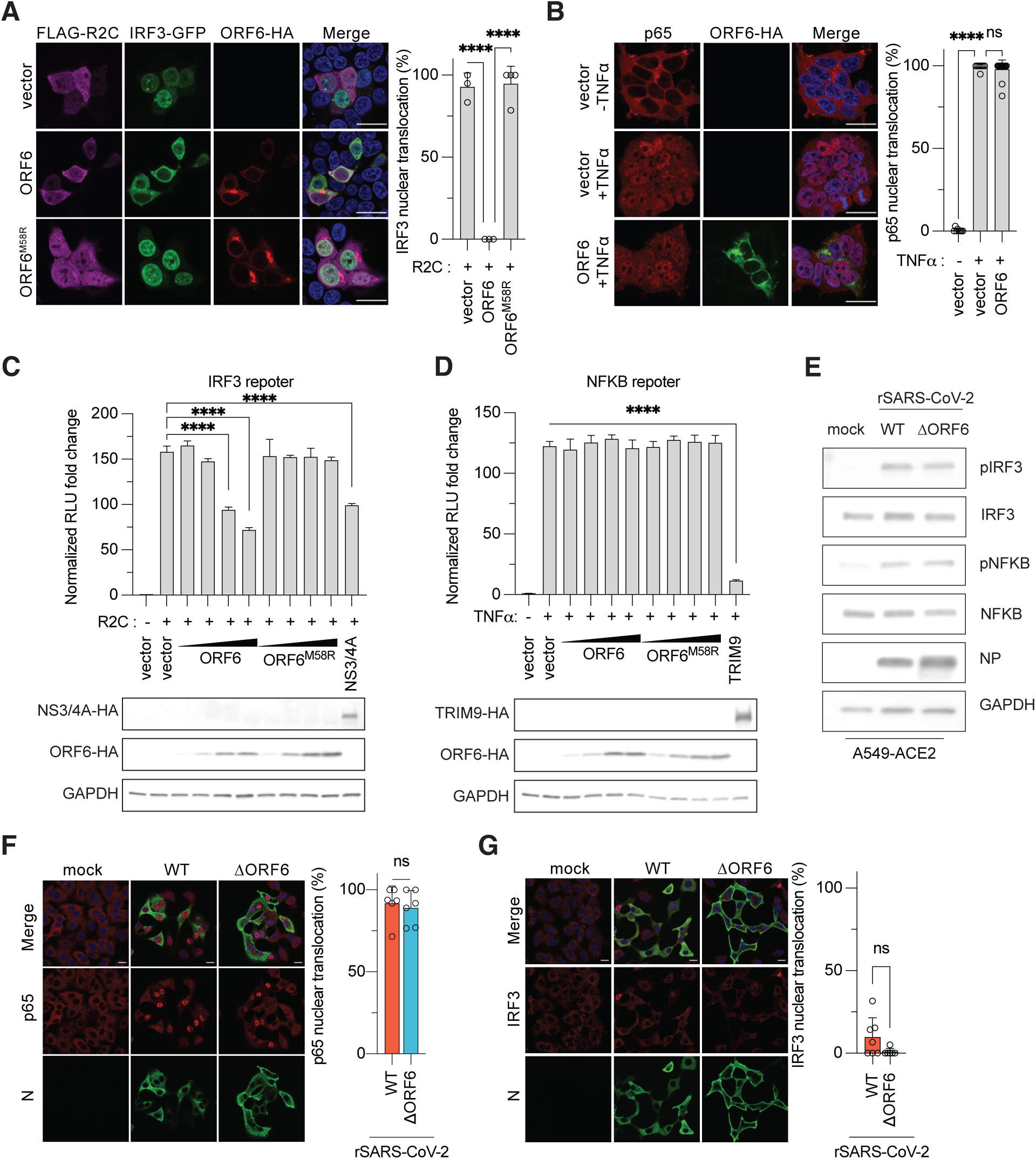
ORF6 selectively blocks nuclear import of innate immune transcription factors. (A) Confocal microscopy images of HEK293T cells transfected with SARS-CoV-2 ORF6, ORF6-M58R or empty vector along with FLAG-RIG-I-2CARD and IRF3-GFP. At 24 h post-transfection, cells were fixed and processed for assessment of the subcellular localization of IRF3-GFP by confocal microscopy. Nuclear translocation of IRF3-GFP in control and ORF6/RIG-I-2CARD double-positive cells was quantified from three fields of view collected from two independent experiments. Data are shown as average ± SD. (B) Confocal microscopy images of HEK293T cells transfected with SARS-CoV-2 ORF6, ORF6-M58R or empty vector. At 24 h post-transfection, cells were treated with TNF-a (25ng/mL) for 45 min. Cells were fixed and processed for assessment of the subcellular localization of p65 by confocal microscopy. Nuclear translocation of p65 in control and ORF6-positive cells was quantified from four fields of view collected from two independent experiments. Data are shown as average ± SD. (C) HEK293T cells were transiently transfected with plasmids expressing ORF6 or ORF6-M58R (0.5 ng, 2 ng, 5 ng, or 10 ng) or HCV NS3/4A (50ng), FLAG-RIG-I-2CARD (5 ng), a plasmid encoding an 3xIRF3-firefly luciferase reporter (p55C1-luc), and plasmid expressing Renilla luciferase from the TK promoter. Data are representative of three independent experiments and shown as average ± SD (n = 3). Cell lysates from the reporter assay were analyzed by Western blot to show relative expression of each transfected viral protein. GAPDH was used as loading control. (D) HEK293T cells were transiently transfected with plasmids expressing ORF6, ORF6-M58R (0.5 ng, 2 ng, 5 ng, or 10 ng), or TRIM9 (100ng), a plasmid encoding an NFKB-firefly luciferase reporter, and plasmid expressing Renilla luciferase from the TK promoter. At 24h post-transfection, cells were treated with 25 ng/mL of TNF-a for 16h, lysed and used for dual luciferase reporter assay. Data are representative of three independent experiments and shown as average ± SD (n = 3). Cell lysates from the reporter assay were analyzed by Western blot to show relative expression of each transfected viral protein. GAPDH was used as loading control. (E) A549-ACE2 cells were either mock-infected or infected with the indicated recombinant viruses at MOI 0.5 for 24 hours. Expression and phosphorylation status of the of the indicated proteins was determined by Western Blot using GAPDH as a loading control. (F) A549-ACE2 cells were infected with the indicated viruses at MOI 0.5. At 24 h post-transfection, cells were fixed and processed for assessment of the subcellular localization of p65 by confocal microscopy. Nuclei were stained with DAPI. IRF3 nuclear translocation in infected and bystander cells was quantified from ≥150 cells per condition from two biological replicates and compared to translocation in mock-infected cells. Data are shown as average ± SD. (G) Same as F but subcellular localization of IRF3 was assessed by confocal microscopy. (Scale bar = 20 µm).). Data in A-D were analyzed by ordinary one-way ANOVA using Turkey’s multiple comparison test. Data in E-F were analyzed by two-tailed unpaired Students T-test. P > 0.05 = ns, P < 0.0001 = ****. Graphs were generated with Graphpad PRISM (version 9).

### SARS-CoV-2 ORF6 disrupts mRNA nuclear export and inhibits host gene expression

As we and others have previously shown that SARS-CoV-2 infection results in the inhibition of host mRNA nuclear export (*11, 12*), we sought to investigate whether the ORF6-Nup98/Rae1 interaction could contribute to this process. To this end, we first transfected HEK293T cells with plasmids encoding SARS-CoV-2 ORF6, ORF6^M58R^ or empty vector, and looked at the intracellular distribution of bulk Poly(A) RNA levels by fluorescence in situ hybridization (RNA-FISH). Remarkably, while bulk Poly(A) RNA was localized throughout the cell in empty vector transfected cells, expression of wildtype ORF6, but not ORF6^M58R^, resulted in a significant increase in the nuclear to cytoplasmic ratio (N/C) of Poly(A) RNA (Fig. 3A), indicating that ORF6 also disrupts Nup98/Rae1 mRNA nuclear export functions. Next, to further address the contribution of ORF6 to the inhibition of mRNA export during infection, we infected A549-ACE2 cells with rSARS-CoV-2 WT, rSARS-CoV-2 ΔORF6 or rSARS-CoV-2 ORF6^M58R^ and performed nucleocytoplasmic fractionation to quantify the abundance of a set of transcripts previously reported to be retained into the nucleus of SARS-CoV-2 infected cells (*12*) within the two different fractions. Strikingly, in agreement with our RNA-FISH data, when we looked at mRNA N/C ratios we found a significant reduction of nuclear mRNA retention in cells infected with both the ORF6-deleted and ORF6^M58R^ virus as compared to wildtype (Fig. 3B), strongly indicating that ORF6 contributes to the disruption of mRNA export and is likely to inhibit host gene expression during infection.

**Figure 3.**
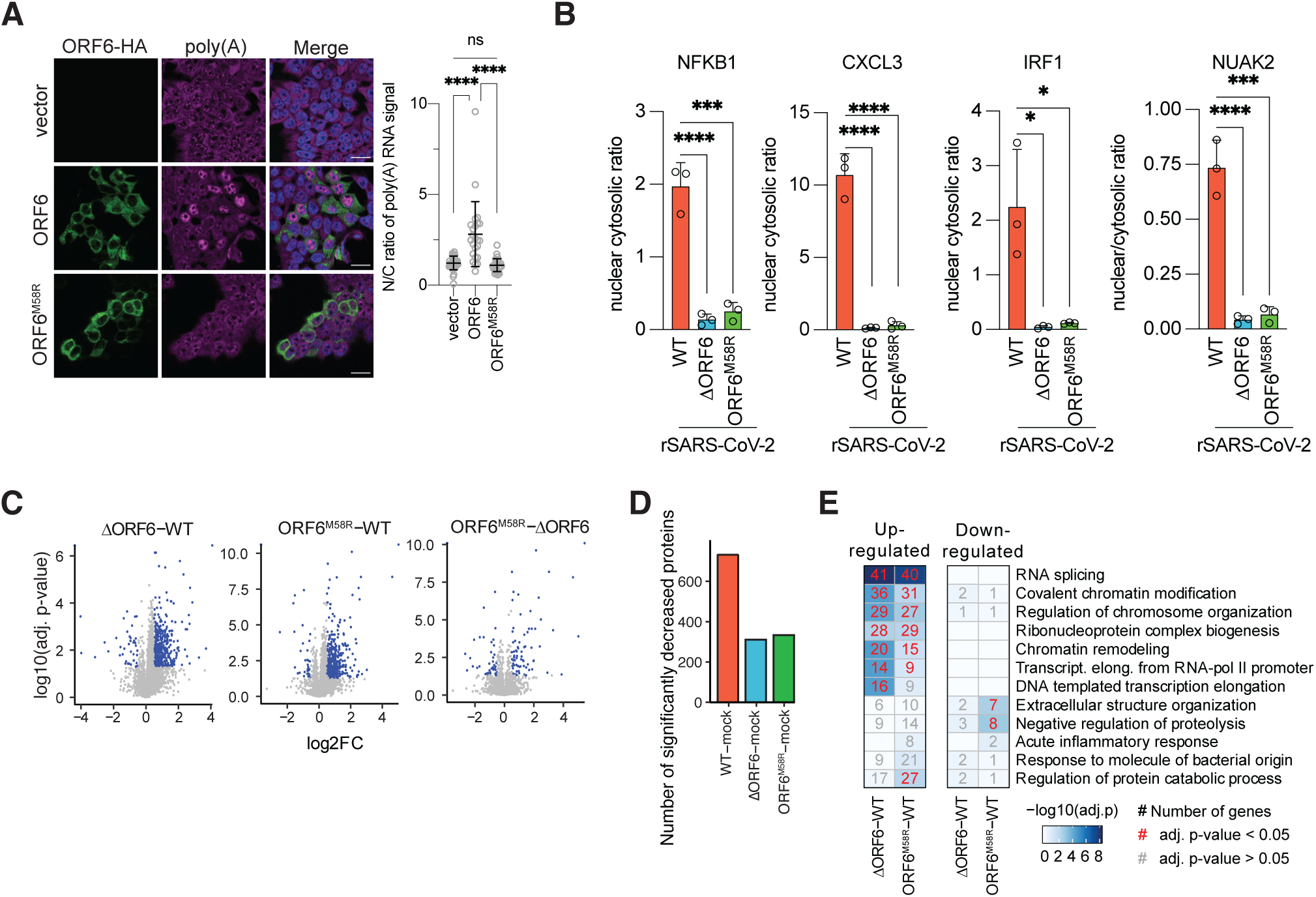
ORF6 disrupts mRNA nuclear export and contributes to host translational shutdown during infection. (A) Confocal microscopy images of HEK293T cells transfected with either empty vector or plasmids encoding ORF6-HA or ORF6 M58R-HA. 24h post-transfection cells were subjected to RNA-FISH analysis as described in methods to detect Poly(A) RNA, and HA immunofluorescence to detect ORF6 expression. The fluorescence intensity of Poly(A) RNA in the nucleus and cytoplasm was determined for ≥ 24 cells per condition and used to compare the ratios of nuclear over cytoplasmic signals for individual cells transfected with either construct (n=2). Data are shown as average ± SD. Significance was determined by unpaired two-tailed t-test: P > 0.05 = n.s., P < 0.0001 = ****. (B) A549-ACE2 cells were infected with the indicated viruses at MOI 2 for 24 hours. Cells were then subject to nuclear cytoplasmic fractionation and RNA was isolated and subjected to RT-qPCR for indicated transcripts. Graph shows nuclear/cytoplasmic ratio of indicated transcripts after normalization to respective compartment markers (GAPDH for cytoplasm, MALAT-1 for nucleus) (n=3). Data are shown as average ± SD. Significance was determined by unpaired two-tailed t-test: P < 0.01 = **, P < 0.0001 = ****. (C) A549-ACE-2 cells were infected with SARS-CoV-2 WT, -ΔORF6, or -ORF6 M58R at MOI 2 for 24h before lysis and processing for mass spectrometry proteomics analysis (see Methods). Volcano plots depict changes in gene expression for indicated comparisons (e.g. ΔORF6 -WT indicates log_2_(ΔORF6/WT)), with log2 fold change (log2FC) on the x-axis and Benjamini-Hochberg adjusted p-values on the y-axis. (D) The number of significantly decreased proteins of each virus compared to mock, based on an absolute value log2FC > 1 and adjusted p < 0.05 cutoff. (E) GO Biological Process enrichment analysis of significantly up- or down-regulated proteins (blue dots in C). Numbers in squares indicate the number of proteins mapping to each term, red numbers indicate a significant enrichment, and grey numbers indicate a non-significant enrichment based on an adjusted p-value < 0.05 cutoff. Additionally, heatmap colors map to the -log_10_ adjusted p-values. (Scale bar = 20 µm). Data in A-B were analyzed by ordinary one-way ANOVA using Turkey’s multiple comparison test. Data in C-E were analyzed as described in methods. P > 0.05 = ns, P < 0.05 = *, P < 0.001 = ***. P < 0.0001 = ****. Graphs were generated with PRISM (version 9).

To further explore the global effect of ORF6 on host gene expression, we next performed mass spectrometry abundance proteomics and phosphoproteomics (Fig. 3 and S2A-E). Importantly, in order to ensure comparable infection rates, in these experiments A549-ACE2 cells were infected with the three recombinant viruses at an MOI of 2 for 24 hours, resulting in comparable infections rates (Fig. S2A). As expected, principal component analysis (PCA) of the abundance proteomics data showed that infected cells clustered away from uninfected cells along the first principal component, suggesting a shift in protein expression upon infection (Fig. S2B). In addition, cells infected with the ORF6-deficient virus clustered together with the rSARS-CoV-2 ORF6^M58R^ infected samples, suggesting that ORF6 expression dramatically remodels host gene expression primarily by altering Nup98/Rae1 nuclear transport functions. In line with the observed ORF6 - mediated disruption of nucleocytoplasmic trafficking, we found that cells infected with the ORF6-deficent or ORF6^M58R^ viruses showed an overall increase in host protein expression with respect to cells infected with rSARS-CoV-2 WT, while a comparison between the ORF6-mutant viruses indicated a more similar protein expression profile (Fig. 3C-D). Interestingly, gene ontology (GO) analysis revealed that the top biological processes upregulated during infection with the ORF6 mutant viruses are linked to mRNA metabolism and include RNA splicing, ribonucleoprotein biogenesis, RNA polymerase II elongation, among others (Fig. 3E). Similar GO biological processes also appear to be regulated by ORF6 at the level of protein phosphorylation (Fig. S2D).

### ORF6 modulates viral protein expression

Based on the observation that SARS-CoV-2 N protein expression was consistently upregulated in cells infected with the ORF6-deficient as compared to the wildtype virus throughout our experiments (Fig. 1D-E, 2E, and S2A), we hypothesized that ORF6 could play a role in the modulation of viral gene expression. To test this hypothesis, we first quantified the differences in viral protein levels between rSARS-CoV-2 WT and rSARS-CoV-2 ΔORF6 infected cells from our global abundance proteomics. As shown in Fig. 4, our analysis revealed significant differences in the relative abundance of several ORF1a/b- and subgenomic RNA (sgRNA)-derived viral proteins, despite very similar infection rates (Fig. S2A). Remarkably, we found that the levels of all ORF1ab-derived NSPs, with the exception of NSP15 and NSP16, were significantly down-regulated in rSARS-CoV-2 ΔORF6 infected cells. However, expression of several structural and accessory proteins, namely S, ORF3A, M, ORF7A, N and ORF9B, was up-regulated compared to wildtype virus infected cells (Fig. 4A). This phenotype of reduced NSP levels and increased expression of sgRNA-derived viral proteins by the ORF6-deficient virus was also validated by Western blot analysis of A549-ACE2 cells lysed 24 hours post-infection (Fig. 4B).

**Figure 4.**
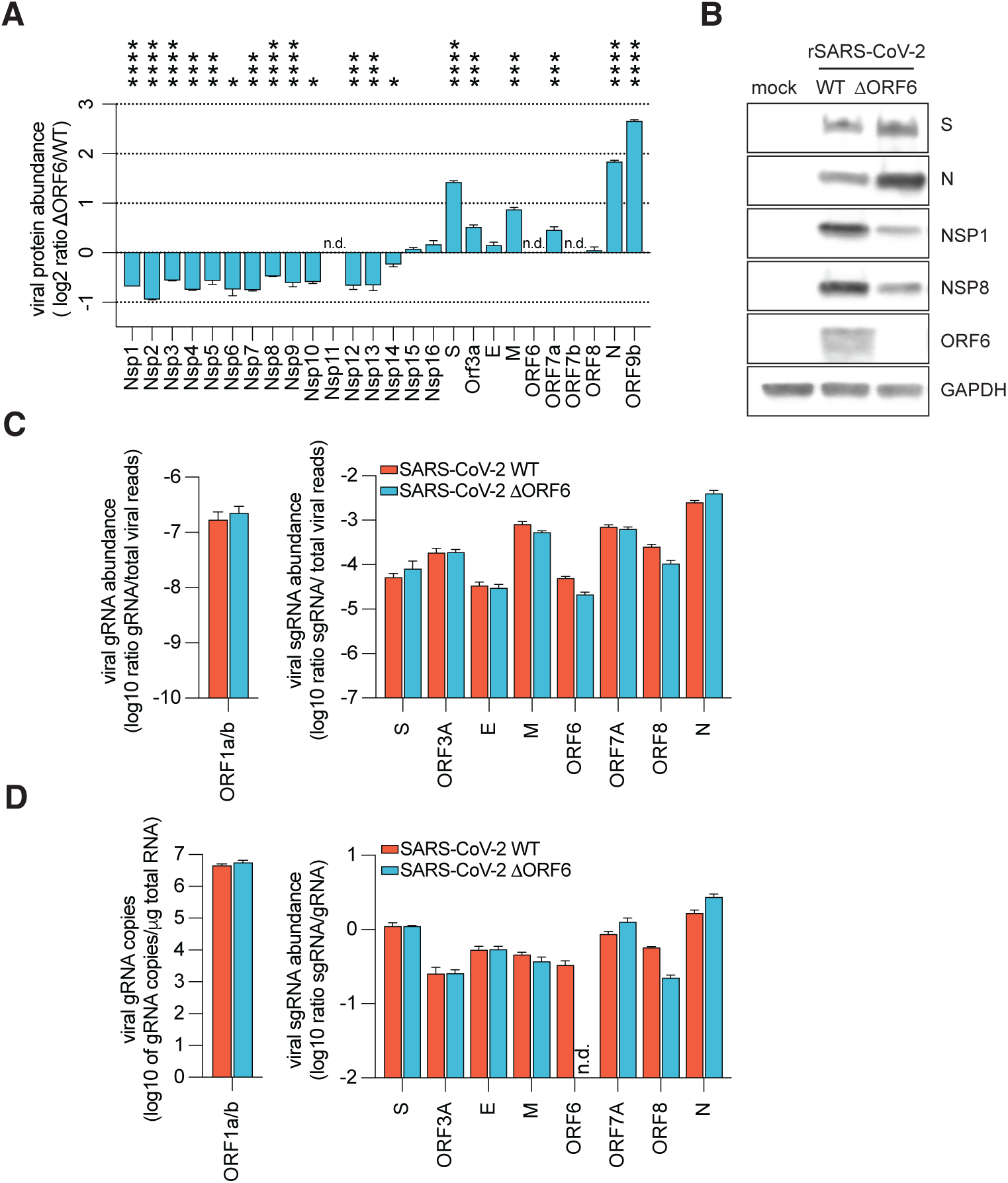
Comparison of viral RNA and protein expression between rSARS-CoV-2 WT and rSARS-CoV-2 ΔORF6. (A) Expression of viral proteins from mass spectrometry abundance proteomics. A549-ACE2 cells were infected with rSARS-CoV-2 WT or ΔORF6 at MOI 2 (n=3 for wt, n=2 for ΔORF6). Graph shows log2 ratio of summed peptide intensities per viral protein of ΔORF6-infected over WT-infected cells. Data are shown as average ± SD. (B) Abundance of the indicated viral proteins was assessed by Western blot from lysates described in A using GAPDH as loading control. (C) Viral gRNA and sgRNA abundance in A548-ACE2 cells infected with indicated viruses at MOI 0.5 for 24h (n=3). Data are shown as ratio of mapped reads of indicated viral RNA species over the sum of viral reads per sample. (D) Viral gRNA copy number per µg of total RNA and ratio of indicated sgRNAs over gRNA per sample from samples described in C as determined by qRT-PCR (n=3). Data in A-B were analyzed by two-tailed unpaired Students multiple T-test. Data in C-D were analyzed by Man-Whitney test with an false detection rate of 5%. Graphs were generated with PRISM (version 9).

Next, to investigate whether ORF6 modulates viral protein expression at a transcriptional or post-transcriptional (translation and/or protein stability) step, we infected A549-ACE2 cells with either rSARS-CoV-2 WT or rSARS-CoV-2 ΔORF6 and quantified the levels of sgRNA/total viral reads and sgRNA/gRNA by bulk mRNA sequencing and quantitative real time PCR (qRT-PCR), respectively (Fig. 4C-D). As shown in Fig. S2F, similar infection rates were achieved by the two viruses under these experimental conditions. Interestingly, we found that despite the remarkable modulation of viral protein expression, the production of both genomic and subgenomic transcripts was only marginally affected in cells infected with the ORF6-deficient virus (Fig. 4C-D). Thus, these results suggest that ORF6 plays a previously unrecognized role in the SARS-CoV-2 life cycle and is critical for the post-transcriptional modulation of viral protein expression. Importantly, while the molecular mechanisms underlying this ORF6 function are still under investigation, the similar levels of N protein expression observed in rSARS-CoV-2 WT and rSARS-CoV-2 ORF6^M58R^ infected cells (Fig. 1E and S2A), strongly suggest that this process is independent of the interaction of ORF6 with the Nup98-Rae1 complex.

### ORF6 expression contributes to viral pathogenicity in Syrian golden hamsters

Next, to evaluate the role of ORF6 in the pathogenesis of SARS-CoV-2, 8-week-old female hamsters were either mock-infected or intranasally inoculated with 5×10^5 PFU of either the parental rSARS-CoV-2 WT or the ORF6-deficient virus. Daily weights were recorded in the 3 groups from the day of infection up until 15 days post-infection (dpi). In addition, animals from each group were sacrificed at 2, 4, and 6 dpi and lungs and nasal turbinates were collected and processed for viral titer determination and histopathological evaluation (Fig. 5A). Remarkably, we found that animals infected with the ORF6 deficient virus exhibited significantly reduced body weight loss and began to recover approximately 3 days earlier than animals infected with the wildtype-virus (Fig. 5B). However, we did not find significant differences in viral titers in both lung and nasal turbinates (Fig. 5C). These results suggest that rather than the viral load, changes in the host response to the infection between the two viruses are likely to be responsible for the observed differences in morbidity. Next, to evaluate the impact of viral infection in the lungs of infected animals, we performed a detailed histopathological evaluation on lungs collected at 2, 4, and 6 dpi. Temporal histologic phenotypes observed in rSARS-CoV-2 WT and ORF6 deficient virus were not readably discernible qualitatively and were consistent with previous reports of COVID-19 in Syrian golden hamsters (*26*). In brief, this was characterized by necrosuppurative bronchiolitis at 2 dpi that progressed to bronchointerstitial pneumonia with edema and hemorrhage at 4 dpi, culminating in a reparative response reflected by bronchiolar and alveolar type 2 (AT2) cell hyperplasia and bronchiolization of alveoli at 6 dpi. However, subsequent quantitative tissue classification of H&E-stained lung sections revealed that wildtype-virus infected animals exhibited a significant increase in the percentage of consolidated lung area at 6 dpi compared to animals infected with the ORF6 deficient virus (Fig. 5E). Histologically this was reflected by an increased proliferative index as determined by the percentage of nuclei expressing Ki67, which predominated in areas of AT2 cell hyperplasia (Fig. 5F). Taken together these findings suggest that a more robust reparative response occurs in wildtype infected hamsters attributable to increase lung injury at earlier timepoints that correlates with the difference lung/bodyweight ratio at 6 dpi (Fig 5D).

**Figure 5.**
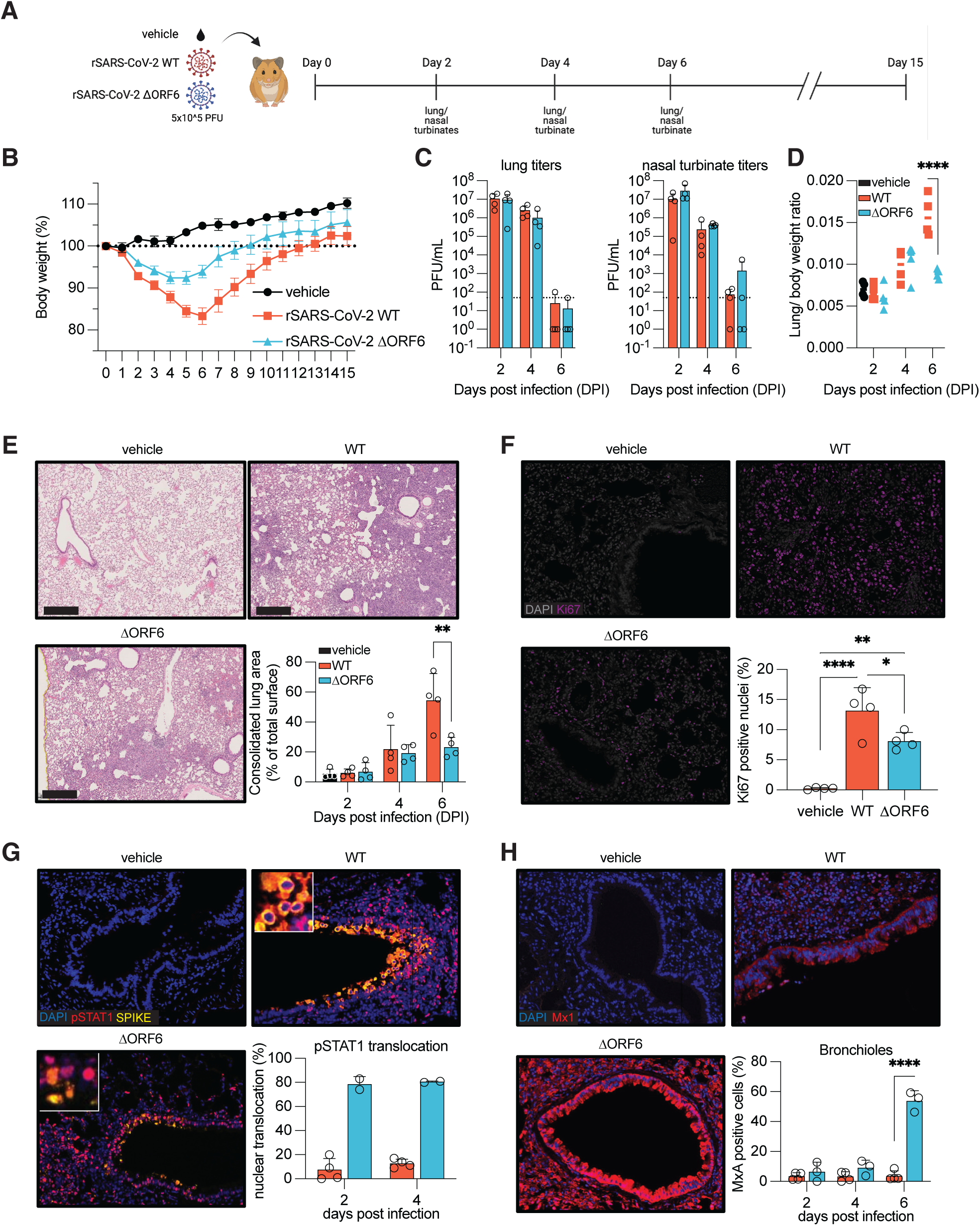
ORF6 plays a critical role in SARS-CoV-2 pathogenesis in Syrian hamsters. (A) Schematic of the *in vivo* experiment using Golden Syrian hamsters that were either mock infected or infected with rSARS-CoV-2 WT or ΔORF6 (n=8 for mock, n=17 for WT and ΔORF6). (B) Hamster weight for the duration of the experiment as a percentage of their weight on day 0. Weight loss data is shown as mean ± SEM. (C) Lung and nasal turbinate viral titers for infected animals at indicated days post-infection. Dashed line indicates the limit of detection for plaque assay (50 PFU/mL) (n=4). (D) Lung to body weight ratio for animals sacrificed at the indicated days post-infection. Line indicates mean value (n=4). (E) Representative images of lung H&E staining for all three groups of animals at day 6 post-infection. Graph shows the consolidated lung area for mock-infected or SARS-CoV-2 WT or ΔORF6 -infected animals at the indicated time points. Data are shown as average ± SD (n=4). Scale bars in histology slides = 500 μm. (F) Tissue sections from mock-, WT-, or ΔORF6-infected animals from 6 days post-infection were stained for DAPI and Ki67 (n=4). Ki67-positive nuclei were quantified as described in methods. Data are shown as average ± SD. (G) Tissue sections from mock-, WT-, or ΔORF6-infected animals were stained for DAPI, pSTAT1, and SARS-CoV-2 Spike (n=3 for WT, n=2 for ΔORF6). pSTAT1 nuclear translocation was quantified as described in methods. Data are shown as average ± SD. (H) Tissue sections from mock-, WT-, or ΔORF6-infected animals were stained for DAPI and Mx1 (n=3). Mx1 expression was quantified as described in methods. Data are shown as average ± SD. Data in C-E and H were analyzed by two-way ANOVA using Šídák’s multiple comparisons test. Data in F were analyzed by ordinary one-way ANOVA using Turkey’s multiple comparison test. P < 0.05 = *, P < 0.01 = **. P < 0.0001 = ****. Graphs were generated with PRISM (version 9).

Given the prominent role of ORF6 in the inhibition of IFN signaling, we next sought to assess STAT1 nuclear translocation in infected cells by performing immunohistochemistry (IHC) for SARS-CoV-2 S and pSTAT1 in lungs harvested at 2 and 4 dpi. In agreement with our previous findings *in vitro*, we found that pSTAT1 was mainly localized in the cytoplasm of S positive cells within the bronchioles of rSARS-CoV-2 wildtype infected animals. However, approximately 80% of the double positive cells in the lungs of animals infected with the ORF6 deficient virus showed a nuclear pSTAT1 staining at both 2 and 4 dpi (Fig. 5G). This is also consistent with the ability of ORF6 to inhibit IFN-mediated STAT nuclear translocation in a Syrian golden hamster cells line (BHK-21) that we observed upon ectopic expression (Fig. S3). Importantly, the impact of ORF6 in the inhibition of IFN signaling was also corroborated by the augmented Mx1 protein expression detected by IHC at 6 dpi in the lungs of the rSARS-CoV-2 ΔORF6 infected animals (Fig. 5H).

Lastly, since infection with SARS-CoV-2 is not lethal in the Syrian golden hamster model, we evaluated the ability of the rSARS-CoV-2 ΔORF6 virus to trigger a protective immune response against a challenge with the wildtype virus. As shown in Fig. S4A, at 30 days after the initial infection, 4 animals from each group were re-challenged with 1×10^5 PFU of SARS-CoV-2 WA/01 and monitored daily for body weight loss. As expected, infection of mock-treated hamsters led to approximately 20% body weight loss by 6 dpi. However, no changes in body weight were observed in hamsters previously infected with the rSARS-CoV-2 wildtype or rSARS-CoV-2 ΔORF6 virus (Fig. S4B). Consistently, similar levels of serum immunoglobulin G (IgG) against full-length viral S protein (Fig. S4C), and no infectious virus in the nasal washes (Fig. S4D), were detected in hamsters previously challenged with the recombinant wildtype and ORF6-deficient viruses.

### The ORF6^D61L^ mutation shared by Omicron variants BA.2 and BA.4 disrupts protein functions at the NPC

Despite the sporadic emergence of frameshifts and/or nonsense mutations in ORF6 during the current COVID-19 pandemic, such mutations have not spread dominantly in the viral population until recently (*27*). We previously reported that the upregulation of key viral innate antagonists, including ORF6 and ORF9b, by the Alpha variant of concern (VOC) likely contributed to its enhanced transmission and human adaptation (*20*). Interestingly, a single point mutation in ORF6 (ORF6^D61L^) recently emerged in the Omicron subvariants BA.2 and BA.4 that is no longer present in the now dominant BA.5 subvariant, which otherwise shares a lot of similarities with BA.4 (*19*). The ORF6 D61 residue is located in close proximity to the key M58 residue at the C-terminal tail (CTT) of the protein that directly binds to the RNA binding pocket of the Nup98-Rae1 complex (*15, 28*) and accompanying paper). Therefore, we sought to investigate the impact of this mutation on the ability of ORF6 to interact with the Nup98-Rae1 complex and inhibit IFN signaling. To this end, we ectopically expressed HA-tagged ORF6, ORF6^D61L^, ORF6^M58R^, or empty vector in HEK293T cells and assess their ability to pull-down endogenous Nup98-Rae1 from cell lysates upon HA immunoprecipitation (Fig. 6A). As expected, we found that ORF6 binds to both Nup98 and Rae1, and that the M58R mutation is sufficient to abolish this interaction (*8*). Strikingly, binding of ORF6^D61L^ to Nup98 and Rae1 was significantly reduced, indicating that the D61 residue is important for interaction with the NPC (see also accompanying paper).

**Figure 6.**
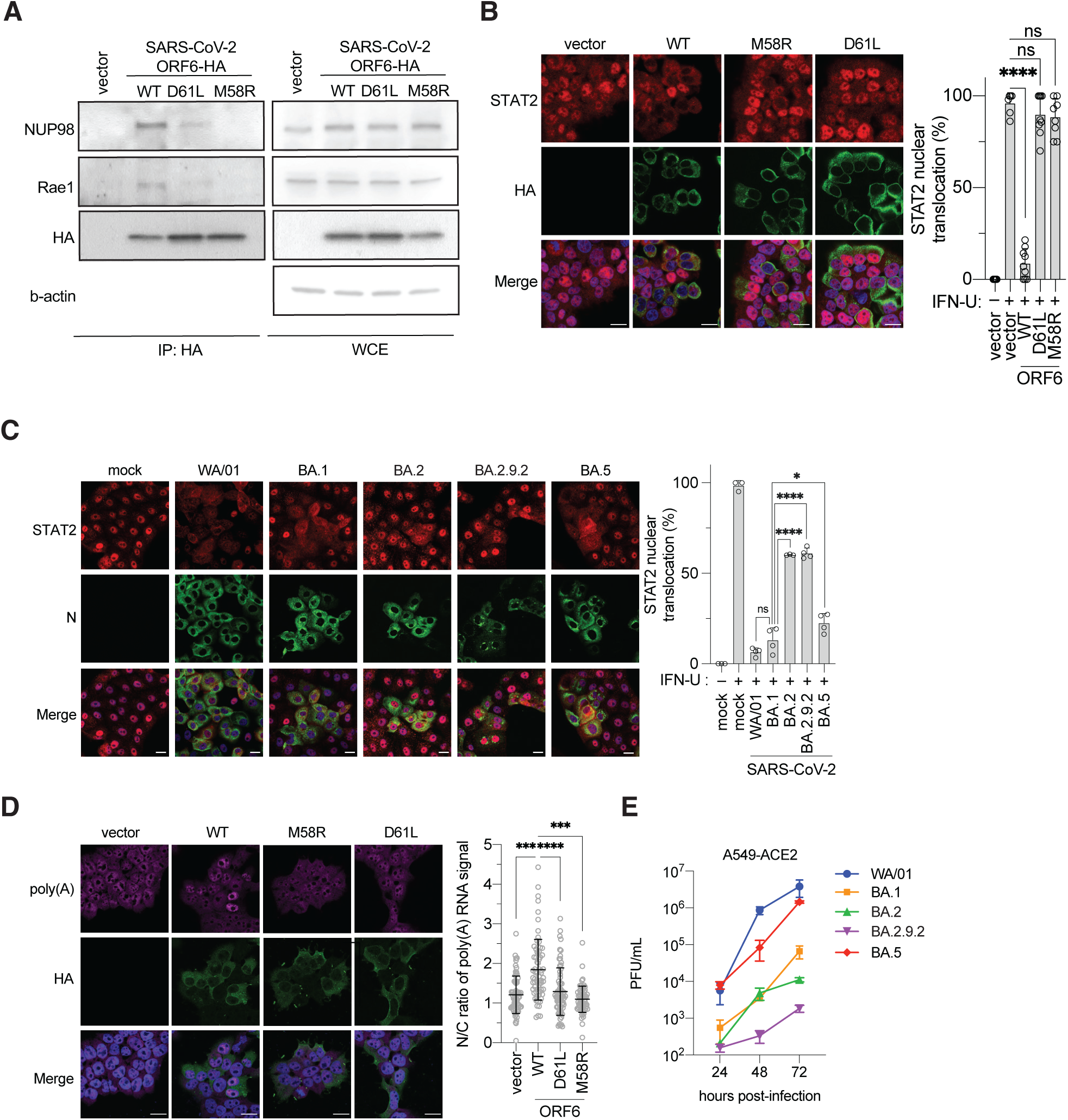
Characterization of the ORF6 D61L mutation. (A) HEK293T cells were transfected with empty vector a construct encoding ORF6, ORF6 D61L, or ORF6 M58R for 24h before lysis. Next, lysates were subjected to HA-tag immuno-precipitation as described in methods before analysis by Western blot for interaction with indicated proteins (IP:HA = eluate after immunoprecipitation, WCE = whole cell extract) (n=3). (B) Confocal microscopy images of HEK293T cells transfected with either empty vector or plasmids encoding ORF6, ORF6-D61L, or ORF6 M58R. At 24h post-transfection cells were treated with 1000 U of IFN universal for 45 min prior to fixation. The subcellular localization of STAT2 was analyzed by confocal microscopy. Nuclei were stained with DAPI. STAT2 nuclear translocation was quantified from ≥150 cells per condition from two biological replicates and compared to translocation in mock-infected cells. (C) Confocal microscopy images of HEK293T cells transfected with either empty vector or plasmids encoding ORF6-HA, ORF6-D61L ORF6 M58R-HA. 24h post-transfection cells were subjected to RNA-FISH analysis as described in methods to detect Poly(A) RNA and HA immunofluorescence to detect ORF6 expression. The fluorescence intensity of Poly(A) RNA in the nucleus and cytoplasm was determined for 50 cells per condition used to compare the ratios of nuclear over cytoplasmic signals for individual cells transfected with either construct. Data are shown as average ± SD. (D) Confocal microscopy images of A549-ACE2 cells infected with the indicated viruses at MOI 0.5 for 24 hours prior to fixation. The subcellular localization of STAT2 was analyzed by confocal microscopy. Nuclei were stained with DAPI. STAT2 nuclear translocation was quantified from ≥150 cells per condition from two biological replicates and compared to translocation in mock-infected cells. Data are shown as average ± SD. (E) Growth curve of indicated viruses at 24h, 48h, and 72h post-infection in A549-ACE2 cells after initial infection at MOI 0.1. Titers were quantified by plaque assay in TMPRSS2-Vero E6 cells. (Scale bar = 20 µm) Data in B-D were analyzed by ordinary one-way ANOVA using Turkey’s multiple comparison test. Graphs were generated with PRISM (version 9). P > 0.05 = ns, P < 0.05 = *, P < 0.001 = ***. P < 0.0001 = ****. Graphs were generated with PRISM (version 9).

Next, we examined the impact of the ORF6^D61L^ mutation on nuclear-cytoplasmic trafficking. Remarkably, we found that ectopic expression of ORF6^D61L^ was unable to block IFN- and 2CARD-RIGI-mediated nuclear translocation of STAT2 and IRF3, respectively (Fig. 6B and Fig. S5A-B). In addition, STAT2- and IRF3-dependent gene expression was also not affected by ORF6^D61L^ overexpression in our luciferase-based reporter assays (Fig. S5C-E). In agreement with these findings, we observed a significant increase in the levels of IFN-dependent STAT2 nuclear translocation in cells infected with Omicron subvariants carrying the ORF6^D61L^ mutation as compared to cells infected with the WA/01 ancestral strain or the Omicron BA.1 and BA.5 VOCs (Fig. 6C). Importantly, we also looked at the intracellular distribution of bulk Poly(A) RNA levels by RNA-FISH and found that ORF6^D61L^, unlike wildtype ORF6, did not significantly increase the N/C ratio of poly(A) RNA in the cell. This indicates that the D61L mutation is also able to interfere the ability of ORF6 to disrupt Nup98/Rae1 mRNA nuclear export functions (Fig. 6D). Because of the significant impairment of ORF6 functions at the NPC by the D61L mutation, we then tested whether the replication of Omicron BA.2 and BA.2.9.2, both expressing ORF6^D61L^, was attenuated in IFN-competent A549-ACE2 cells. Intriguingly, as shown in Fig. 6E, we found that these two subvariants replicate less efficiently as compared to BA.1 and BA.5. All together these results suggest that the D61L mutation significantly disrupts ORF6 protein functions at the NPC and impairs innate immune evasion with potential implications for viral fitness.

## DISCUSSION

The innate immune response acts as a first line of defense against infection by upregulating IFN-stimulated genes (ISGs) expression and limiting virtually any step of the virus life cycle to promote viral clearance(*29*). As a countermeasure, SARS-CoV-2 has evolved multiple strategies to suppress or at least interfere with the IFN response and enhance replication and transmission (*1, 7, 20, 30*). In this study, we used molecular and biochemical methods, combined with *in vivo* animal studies, to dissect the impact of the innate immune antagonist ORF6 on the host response to SARS-CoV-2 infection. Our results show that ORF6, by interacting with the Nup98-Rae1 complex at the nuclear pore, can interfere with nucleocytoplasmic trafficking in two distinct ways: by selectively inhibiting karyopherin-mediated nuclear import pathways and by modulating host cell mRNA export.

Using recombinant wildtype and ORF6-mutant viruses as well as recently emerged SARS-CoV-2 Omicron variants containing a D61L mutation in the C-terminal tail of the protein, we show that ORF6-Nup98 interaction is required to block STAT1 and STAT2 nuclear translocation during infection, thereby inhibiting ISG expression, both *in vitro* and in the Syrian golden hamster model *in vivo.* In addition, we also found that ORF6 cannot prevent nuclear translocation of NF-kB p65, which has also been shown to be mediated by the classic karyopherin alpha/beta pathway (*31*), pointing towards a selective inhibition of nuclear import. Interestingly, while it cannot be excluded that some cargos could traffic using an alternate route if the karyopherin alpha/beta pathway is blocked, such specificity may also suggest the existence of different subsets of Nup98-dependent and Nup98-independent cargo complexes. Further studies will be required to fully understand the molecular basis for this specificity.

A second common mechanism to inhibit host gene expression and downregulate innate antiviral defenses is to interfere with nuclear mRNA export (*32–34*), and different viruses have been shown to target the Nup98-Rae1 complex to accomplish this effect (*35–38*). We previously showed that SARS-CoV-2 infection inhibits host mRNA nuclear export and that the viral NSP1 protein contributes to this process by binding to the mRNA export receptor heterodimer NXF1-NXT1 and reducing its interaction with the NPC (*11*). Since NXF1-NXT1 interacts with phenylalanine-glycine (FG) repeats on nucleoporins, such as Nup98, to mediate docking of messenger ribonucleoproteins (mRNPs) and facilitate trafficking trough the NPC (*39, 40*), we hypothesized that ORF6 could also interfere with this process. In addition, structural data have recently shown that, similarly to VSV M and herpesvirus ORF10 proteins (*37, 38*), the CTT of ORF6 directly interacts with the RNA binding groove of the Nup98-Rae1 complex and competes for in vitro binding of single-stranded RNA (*15, 28*). Consistent with these notions, our results show that ORF6 expression can indeed block Nup98-Rae1 mRNA export functions and this contributes to the shutoff in protein synthesis that occurs during SARS-CoV-2 infection. Furthermore, we observed that the D61L mutation, shared by the Omicron variants BA.2 and BA.4, in addition to interfere with the ability of ORF6 to inhibit nuclear import, by disrupting ORF6 binding to Nup98 - Rae1 also influences its ability to block host mRNA export, which may have important implications for viral transmissibility and pathogenicity (see accompanying paper). Since NSP1 has also been shown to inhibit host mRNA export, a better understanding of how ORF6 and NSP1 functions cooperate and/or complement each other during infection will be key to fully reveal the molecular mechanisms underlying innate immune antagonism by these viral proteins.

Consistent with a role of ORF6 in viral pathogenesis, hamsters infected with an ORF6 deleted SARS-CoV-2 experienced less body weight loss and reduced lung injury and AT2 cell hyperplasia, correlating with increased STAT2 translocation and ISG expression in lungs. Surprisingly, this did not result in significant lower levels of viral replication in the respiratory tract of hamsters for the ORF6 minus virus. The impact of ORF6 function in viral replication in hamsters might be too subtle to be able to detect in this experimental animal model. A small impact in viral replication *in vivo* associated with ORF6 function might also explain the circulation and transmission of Omicron BA.2 and BA.4 variants containing a deleterious ORF6 polymorphism in humans. Nevertheless, these variants are less replicative in IFN-competent human A549 cells than BA.5, lacking this polymorphism, but containing identical changes in spike associated with immuno-evasion as compared to BA.4. BA.5 human transmission dominance over BA.4 might be at least in part mediated by the lack of the ORF6 D61L mutation (see also accompanying paper).

Interestingly, our work has also revealed a previously unobserved role of ORF6 in the modulation of viral protein expression. Indeed, we discovered that the relative expression of several ORF1a/b- and sgRNA-derived viral proteins was significantly altered in cell infected with an ORF6-deficient SARS-CoV-2 virus, to favor expression of several structural (S, M and N) and accessory viral genes (ORF3A, ORF7A and ORF9B). At this time, it remains unclear whether this phenomenon is mediated by a direct role of ORF6 on the translational or post-translational regulation of viral gene expression, or a consequence of the altered activity of some of the other viral proteins. However, due to the comparable levels of viral protein expression we observed between the wildtype and ORF6^M8R^ virus, this phenotype is likely independent of ORF6 functions related to the Nup98-Rae1 complex.

Overall, our data strongly suggests that ORF6 is a major SARS-CoV-2 innate immune antagonist. We show that the absence of ORF6, or the introduction of ORF6 loss-of-function mutations, significantly influences the host antiviral responses resulting in SARS-CoV-2 attenuation both *in vitro* in IFN-competent cells, and *in vivo* in the hamster model. In addition, we functionally characterized the ORF6^D61L^ mutation shared by the BA.2 and BA.4 Omicron variants that have been recently displaced by the now dominant BA.5 VOC, highlighting the importance of genomic surveillance and variant analysis to better understand the mechanisms underlying SARS-CoV-2 evolution, pathogenicity, and immune evasion strategies.

## ACKNOWLEDGEMENTS

We thank all the members of the A.G.-S. laboratory for helpful feedback, in particular Teresa Aydillo and Michael Schotsaert for training and support. We thank Randy Albrecht for support with the BSL3 facility and procedures at the Icahn School of Medicine at Mount Sinai (ISMMS), the staff at the ISMMS Center for Comparative Medicine and Surgery (CCMS) for support with the ABSL3 facility and Richard Cadagan for excellent technical assistance. We are indebted to the efforts of Mount Sinai Pathogen Surveillance Program (Hala Alshammary, Juan David Ramirez, Radhika Banu, Paras Shrestha, Angela A. Amoako, Aria Rooker, Christian Cognigni, Daniel Floda, Adriana van de Guchte, Zain Khalil, Keith Farrugia, Alberto Paniz-Mondolfi) for collection and sequencing of clinical specimens used to derive viral isolates. Confocal microscope images were taken at the Microscopy Shared Resource Facility at the ISMMS. Animal tissues were processed at the Biorepository and Pathology Core (ISMMS) for tissue processing and histology. Bulk RNA Sequencing sample libraries were prepared at the Center for Advanced Genomics Technology Facility directed by Dr. Robert Sebra at ISMMS. This work utilized instruments acquired from NIH SIG grants (S10OD026983 & S10OD030269). Schemes in Fig. 5A and Fig. S4A were created with BioRender.com.

## Data and materials availability

Further information and requests for resources and reagents should be directed to and will be fulfilled by the corresponding authors L.M. and A.G.-S. The mass spectrometry abundance proteomics and phosphoproteomics data have been deposited to the ProteomeXchange Consortium (http://proteomecentral.proteomexchange.org) via the PRIDE partner repository (*41*) with the dataset identifier PXD036821. Bulk RNA-Seq sequence read data and corresponding SARS-CoV-2 sgmRNA processed data files are accessible at the NCBI Gene Expression Omnibus (GEO), accession number pending. Analysis code is available at GitHub (submission pending). The BAC for generation of recombinant SARS-CoV-2 wildtype and mutant ORF6 viruses and their respective viruses are available at this website: https://www.txbiomed.org/business-development/reverse-genetics-plasmids/.

## MATERIALS AND METHODS

### Cell lines

Vero E6 (ATCC, CRL-1586) and TMPRSS2-Vero E6 (BPS Bioscience Cat# 78081) were maintained in Dulbecco’s modified Eagle’s medium (Corning) supplemented with 10% fetal bovine serum (Peak Serum), 1% non-essential amino acids (Gibco), 1% HEPES (Gibco) and 1% penicillin/streptomycin (Corning) at 37 °C and 5% CO2. HEK293T (ATCC, CRL-3216), A549-ACE2 (previously described in (*42, 43*), and BHK-21 (ATCC, CCL-10) were maintained in Dulbecco’s modified Eagle’s medium (Corning) supplemented with 10% fetal bovine serum (Peak Serum) and penicillin/streptomycin (Corning) at 37 °C and 5% CO2. All cell lines used in this study were regularly screened for Mycoplasma contamination, using the Universal Mycoplasma Detection Kit (ATCC, 30-1012K).

### Plasmids, antibodies and cytokines

The expression vectors for SARS-CoV-2 ORF6 and SARS-CoV-2 ORF6^M58R^ plasmids have been described previously (*8*). Constructs encoding FLAG-RIG-I 2CARD (*44*), IRF3-GFP (*44*), HCV NS3/4A-HA (*45*), STAT1-GFP (*46*), HA-TRIM9 (*25, 47*), pRL-TK, ISG54-firefly (*47*), p55C1-Luc (*48*), pNFkB-Luc have been described elsewhere ((*47*). SARS-CoV-2 ORF6 D61L was cloned into the pCAGGS mammalian expression vector that encodes a carboxyterminal HA-tag using NotI and KpnI restriction sites. Overlap-PCR was used to generate the ORF6 D61L mutant by changing residue 61 from aspartic acid to leucine (GAT to CTC). All expression vectors were confirmed by sanger sequencing and are available upon request. Antibodies used for immunoblotting include: anti-HA-HRP (Cell Signaling, 6E2), anti-SARS-CoV-NP antibody (1C7C7), anti-ORF6 (DA087, MRC PPU Reagents and Services), anti-Rae1 (PA5-93166, Thermo-Fisher), anti-Nup98 (2H10, Abcam), anti-phospho-STAT1 (58D6, Cell Signaling), anti-STAT1 (sc-417, Santa Cruz), anti-phospho-STAT2 (D3P2P, Cell Signaling), anti-STAT2 (sc-476, Santa Cruz), anti-GAPDH-HRP (3683S, Cell Signaling), anti-phospho-IRF3 (4D4G, Cell Signaling), anti-IRF3 (D614C,Cell Signaling), anti-phospho-p65 (93H1, Cell Signaling), anti-p65 (D14E12, Cell Signaling), anti-SARS-CoV-2 Spike (2B3), anti-SARS-CoV-2 Nsp1 (GTX135612, Genetex), and anti-SARS-CoV-2 Nsp8 (5A10, Genetex). Secondary antibodies used were anti-mouse-HRP (Kwik, Cat# 1005), anti-rabbit-HRP (Kwik, Cat# 1006), anti-rat-HRP (Invitrogen, #31470), anti-sheep-HRP (A16041, Invitrogen). Antibodies used for immunofluorescence staining are: anti-STAT2 (sc-476, Santa Cruz), anti-SARS-CoV-NP antibody (1C7C7), M2 anti-Flag (Sigma Aldrich, F1804), anti-HA (6E2, Cell Signaling; C29F4, Cell Signaling), anti-phospho-IRF3 (4D4G, Cell Signaling), anti-p65 (D14E12, Cell Signaling), anti-phospho-STAT1 (58D6, Cell Signaling). Secondary antibodies used are: anti-mouse-AlexaFluor488 (Invitrogen, A21202), anti-rabbit-AlexaFluor594 (Invitrogen, A21207), anti-mouse-AlexaFluor647 (Invitrogen, A31571), anti-rabbit-AlexaFluor594 (Invitrogen, A21207), and DAPI (Sigma-Aldrich). Antibodies used for immunostaining of plaque assays are: monoclonal anti-SARS-CoV-NP antibody (1C7C7) and anti-mouse HRP antibody (Abcam ab6823). Antibodies used for immunostaining of histology slides are: SARS-CoV-2 spike (99423S, Cell Signaling) at a 1:400 dilution, goat anti-rabbit HRP-polymer antibody (Vector Laboratories, Burlingame, CA), Ki67 (Dako M061601-2) at a 1:100 dilution, goat anti-mouse HRP-polymer antibody (Vector Laboratories), phospho-STAT1 (9167S, Cell Signaling) at a 1:300 dilution, anti-MxA (EMD Millipore MABF938) at a 1:200 dilution, and DAPI (Akoya Biosciences). The antibody used for used for ELISA is an anti-hamster IgG horseradish peroxidase antibody (HRP, abcam, #ab6892). Cytokines used in this study were Universal Type I Interferon Alpha (PBL Assay Science, cat# 11200-2) and TNF-alpha (Bio-Techne, R&D systems).

### Viruses and infections

All SARS-CoV-2 infections were performed under BSL3 containment in accordance with the biosafety protocols developed by the Icahn School of Medicine at Mount Sinai (ISMMS). Virus infections were performed using SARS-CoV-2, isolate USA-WA1/2020 (BEI Resources NR-52281), SARS-CoV-2 BA.1 (isolate: PV44488), SARS-CoV-2 BA.2 (isolate: PV56107_P2), SARS-CoV-2 BA.2.9.2 (isolate: PV56159_P2), SARS-CoV-2 BA.4 (BEI Resources NR-56806), SARS-CoV-2 BA.5 (isolate: PV58128). Additionally, three recombinant SARS-CoV-2 (rSARS-CoV-2) viruses, based on the USA-WA1/2020 reference sequence were used. The rSARS-CoV-2 WT and rSARS-CoV-2 ΔORF6 have been previously described (*22*). A recombinat virus with a single amino acid mutation in ORF6 at position 58, rSARS-CoV-2 ORF6^M58R^, was generated for this study. rSARS-CoV-2 ORF6^M58R^, was generated using the same bacterial artificial chromosome (BAC)-based SARS-CoV-2 reverse genetic system previously described (*21*). Briefly, two oligonucleotides were used to introduce the M58R coding change into fragment 1 by site-directed mutagenesis (5’-ctcaattagatgaagagcaaccacgggagattgattaaacg-3’ and 5’-tcatgttcgtttaatcaatctcccgtggttgctcttcatct-3’). The region in the wild-type BAC between the unique restriction sites of BamHI and RsrII was replaced by the one from fragment 1 containing the M58R mutation, and the newly generated BAC was used to produce the rSARS-CoV-2 ORF6^M58R^ virus according to the protocol described previously (*23*). All viral stocks were grown in Vero E6 cells (except for Omicron subvariants, which were grown in Vero-TMPRSS2 cells) as previously described and validated by genome sequencing (*8*). Sequencing was either performed using the MinION sequencer (Oxford Nanopore Technologies) or with the Nextera XT DNA Sample Preparation kit (Illumina) as described elsewhere (*49, 50*). Virus growth media (VGM) was used for all infections: Dulbecco’s modified Eagle’s medium (Corning) supplemented with 2% fetal bovine serum (Peak Serum), 1% non-essential amino acids (Gibco), 1% HEPES (Gibco) and 1% penicillin/streptomycin (Corning) at 37 °C and 5% CO2. Viral stocks for *in vivo* studies were concentrated using Amicon ultra-centrifugal filters (100 kDa MW-cutoff, Millipore). All in vivo infections were carried out in a CDC/USDA-approved BSL-3 facility at ISMMS CCMS.

### Plaque assay

Unless otherwise specified, plaque assays were performed using Vero E6 cells in 12-well format as previously described (*51*). Briefly, confluent Vero E6 cells were infected with serial ten-fold dilutions of supernatants of infected cells or supernatants of homogenized tissue. Infections were performed in 12-well format for 1h at 37°C and 5% CO2 using an inoculum of 200uL, rocking plates every 10-15 min. An overlay of MEM with penicillin/streptomycin (Corning), L-Glutamine (Gibco), HEPES (Gibco), BSA (MP Biomedicals), and NaHCO_3_ supplemented with 0.7% purified agar (Oxoid) and 2% fetal bovine serum (Peak Serum) was applied to each well. On day 3 post-infection, cells were fixed with 5% formaldehyde overnight and immuno-stained using a monoclonal anti-SARS-CoV-NP antibody (1C7C7) at a 1:1,000 dilution, an anti-mouse HRP antibody (Abcam ab6823) at a 1:5,000 dilution, and TrueBlue (SeraCare) for detection. All samples were frozen at −80°C once before evaluation of viral titers.

### Western Blot and immunoprecipitation

Vero E6 or A549-ACE2 cells were seeded in a 24-well format at a density of 100,000 cells/well. The next day, cells were infected with SARS-CoV-2 at the indicated MOI in viral growth media for 1 hour after which the inoculum was removed and samples were harvested at 24 hpi. Cells were either lysed directly or stimulated with universal IFN type I (1,000 U/mL) for 45 min before lysis. SARS-CoV-2-infected cells were lysed in radio-immunoprecipitation assay (RIPA) buffer (Sigma-Aldrich) supplemented with 1% sodium dodecyl sulfate, cOmplete protease inhibitor mixture (Roche), and Halt phosphatase inhibitor mixture (Thermo Fisher Scientific) before boiling for virus inactivation. Lysates were normalized for protein concentration using a BCA protein assay (Pierce), supplement with 4X Laemmli sample buffer (Bio-Rad Laboratories), boiled for 10 min, and loaded into 4-20% gradient gels (Bio-Rad Laboratories). Gels were transferred onto polyvinylidene fluoride (PVDF) membranes (Bio-Rad Laboratories) using the Trans-Blot Turbo Transfer System (Bio-Rad Laboratories). Membranes were blocked in Tris-buffered saline with 0.1% Tween 20 detergent (TBS-T) containing 5% nonfat dry milk. Primary antibodies were diluted 1:1,000 in TBS-T containing 3% bovine serum albumin. Secondary horseradish peroxidase-conjugated antibodies were diluted 1:10,000 in TBS-T containing 3% nonfat dry milk. For immunoprecipitation of endogenous Nup98, A549-ACE2 cells were seeded in a 10-cm dish format. Cells were infected with indicated viruses at MOI 2 for 24h. Next, cells were processed as described before (*11*). In brief, cells were lysed in lysis buffer (50 mM Tris (pH 7.5), 150 mM NaCl, 1% IGEPAL CA-630, 0.1 mM Na3VO4, 1 mM NaF, 1 mM DTT, 1 mM EDTA, 1 mM PMSF, 1×cOmplete protease inhibitor mixture and 10% glycerol) for 30 min on ice then incubated at 65°C for 30min to inactivate virus. Inactivated samples were sonicated and then cleared by centrifugation. Lysates were incubated with 10 ug of anti-Nup98 antibody or an irrelevant isotype control (IgG DA1E, Cell signaling) overnight and subsequently incubated with protein G-beads for 2h. Beads were washed and protein was eluted by addition of a 2x sample buffer. Samples were processed following the western blot protocol described above. For immunoprecipitation of ORF6-HA, 500,000 HEK 293T cells were transfected with 1 ug of indicated constructs. At 24 hours post-transfection, cells were lysed in RIPA buffer, cleared by centrifugation, and incubated with EZview Red Anti-HA Affinity Gel beads (Millipore Sigma) at 4 C overnight while shaking. Next, beads were washed for five-times for 5 mins in RIPA buffer at 4 C while shaking before elution of bound proteins by boiling the beads in 2x Laemmli buffer for 10 min at 95 C.

### Luciferase Assay

For luciferase assays, HEK293T cells were seeded in a 24-well format at a density of 100,000 cells/well. The next day, cells were transiently transfected with pRL-TK and either the IRF3 responsive p55C1 promoter (p55C1-Luc) or the NFkB-Luc vector along with the indicated plasmids. For NFkB reporter experiments, cells were treated overnight with human TNF-alpha (25ng/ml) at 24 hours after transient transfection. For the IRF3 reporter experiments, cells were co-transfected with RIG-I-2CARD (5 ng) and lysed at 24 hours after transfection using Passive Lysis Buffer (Promega). Samples were processed and luciferase activity was measured using the Dual-Luciferase Assay System (Promega) according to the manufacturer’s instructions. Firefly luciferase values were normalized to Renilla luciferase values, and the induction was calculated as fold over unstimulated vector control condition.

### Confocal Microscopy

Vero E6, A549-ACE2, HEK293T, or BHK-21 cells were seeded into 24-well glass bottom plates (MatTek) at a low density the day before infection or transfection. For infection experiments, cells were infected at the indicated MOI for 24 hours, then fixed with 5% methanol-free formaldehyde or treated with universal IFN-I at 1,000 U/mL (PBL) before fixation. For overexpression experiments, indicated plasmids were transfected using LT-1 Reagent (Mirus) and cells were then fixed with 5% methanol-free formaldehyde or treated with IFN or TNF-alpha before fixation. IFN treatments were performed for 45 min using universal IFN-I at 1,000 U/mL (PBL). TNF-alpha treatments were performed for 45 min using 25 ng/ml of human TNF-alpha (Thermo Fisher). Cells were permeabilized with 0.1% Triton X in phosphate-buffered saline (PBS) and stained as previously described (*8*). Confocal laser scanning microscopy was performed with a Zeiss LSM880 confocal laser scanning microscope (Carl Zeiss Microimaging) fitted with a Plan Apochromat 63×/1.4 or 40×/1.4 oil objective, or with a 20x/1.4 objective. Images were analyzed with Fiji software (https://fiji.sc/).

### Flow Cytometry

A549-ACE2 cells were seeded in 24-well format at a density of 150,000 cells/well. The next day, cells were infected at indicated MOI for 24 hours. Cells were detached with PBS supplemented with 10 mM EDTA (Gibco) and fixed with 5% formaldehyde. Cells were permeabilized and washed with Perm/Wash buffer (BD), and then stained with monoclonal anti-SARS-NP antibody conjugated to AlexaFluor488 (Invitrogen) for 1 hour. Cells were washed with and resuspended in PBS supplemented with 2% BSA, 2.5 mM EDTA and subsequently subjected to cytometry using a Gallios cytometer (Beckman). 10,000 cells were acquired for each condition. Single cells were gated and the percentage of NP-positive cells was used to determine infection rates for rSARS-CoV-2 WT, ΔORF6, and ORF6^M58R^ viruses. Mean fluorescence intensity of NP-positive cells was also measured for the NP-positive cells in each condition.

### Nuclear-cytosolic fractionation

A549-ACE2 cells were infected at the indicated MOI for 24 hours and subsequently washed with PBS and detached with 10mM EDTA (Gibco). After washing with PBS, cells were resuspended in one volume buffer A (15 mM Tris-HCl pH 8 (Boston Bioproducts), 15 mM NaCl (Corning), 60 mM KCl (Sigma-Aldrich), 1 mM EDTA pH 8 (Invitrogen), 0.5 mM EGTA pH 8 (BioWorld), Spermidine 0.5 mM (Sigma-Aldrich), RNasin 100 U/mL (Thermo-Fisher) and cell membranes were lysed by addition of one volume of buffer A, supplemented with 0.8% NonIdent 40 (US Biological Life Sciences) for 5 min. Cytoplasmic supernatant was separated from nuclei by centrifugation, before washing the nuclei with PBS. Next, nuclei were resuspended in one volume of RLN buffer (50 mM Tris-HCl pH 8, 140 mM NaCl, 1.5 mM Mg2Cl (Sigma-Aldrich), 10 mM EDTA pH 8, RNasin 100 U/mL, 0.8% NonIdent 40) and then lysed by addition of one volume of RLN buffer, supplemented with 0.8% NonIdent 40, for 5 minutes. Debris was removed by centrifugation and cytoplasmic and nuclear fractions were lysed in TriZol reagent (Ambion). 500ng of isolated RNA were reverse transcribed using the High-Capacity cDNA Reverse Transcription Kit (Applied Biosystems) according to manufacturer’s instructions. Quantitative real-time PCR was performed using TaqMan Universal Master Mix II with UNG (Applied Biosystems) according to the manufacturer’s instructions. Cycling program with 50 amplification cycles was designed according to the manufacturer’s instructions. The following TaqMan (ThermoFisher) primer/probe mixes were used: MALAT-1 (Hs00273907_s1), NUAK2 (Hs00388292_m1), NFΚB 1 (Hs00765730_m1), CXCL3 (Hs00171061_m1), IRF1 (Hs00971965_m1), and GAPDH (Hs02786624_g1). Transcripts from each fraction were normalized to a housekeeping gene of the respective compartment (GAPDH for cytosolic fraction, MALAT-1 for nuclear fraction). After normalization, nuclear-cytosolic ratios were calculated for each sample.

### SARS-CoV-2 RT-qPCR

To generate SARS-CoV-2 gRNA standards for quantification of copy numbers, the sequence encoding the section from position 11984 to 13321 in the viral genome, that is covered by the primers used for gRNA amplification (Table 2), was cloned by PCR amplification of viral cDNA into a pGEM vector under control of a T7 promoter using pGEM-T Easy Vector System (Promega). RNA standards were subsequently generated by in vitro transcription using the mMESSAGE mMACHINE™ T7 Transcription Kit (Invitrogen) according to the manufacturer’s instructions. For quantification of viral genome copies during infection, A549-ACE2 were mock-infected or infected at indicated MOI for 24h before lysis in Trizol (Invitrogen). RNA was isolated using DirectZol RNA kit (Zymo Research) or miRNeasy kit (Qiagen) according to the manufacturer’s instructions. Isolated RNA or serial ten-fold dilutions of RNA standards for the ORF1ab amplicon (ranging from 2.25×10^6 to 250 copies/rxn) were reverse transcribed using the Takara Prime Script RT kit (Takara) using poly(A) primers according to manufacturer’s instructions. TaqMan Universal Master Mix II with UNG (Applied Biosystems) was used for the PCR according to the manufacturer’s instructions. Cycling program with 40 amplification cycles was also designed according to the manufacturer’s instructions. GAPDH was used as endogenous gene control and was amplified using the commercial primer/probe set hs02786624_g1 (Applied biosystems). Primers for viral gene amplification were used at 500 nM each, while probes were used at concentration of 250nM. Primer/probe sets were previously described (Table 2) (*52*). For quantification of gRNA and sgRNAs, The LightCycler® 480 SYBR Green I Master mix (Roche) was used according to the manufacturer’s instructions. Cycling program with 50 amplification cycles was also designed according to the manufacturer’s instructions. Primers were used at a final concentration of 1 uM. A leader specific forward primer was used for all reactions and a gene specific reverse primer was designed for each target (Table 2). Results were adjusted for primer efficiency as described previously (*53*).

**Table 1.** Coding changes of viruses used in this study. Changes refer to the reference strain Wuhan-Hu-1.

**TABEL 1.**
| Virus | Coding changes<br>(compared to Wuhan-Hu-1 reference strain) |
| --- | --- |
| SARS-CoV-2 (USA-WA1/2020) | ORF8:L84S |
| recombinant SARS-CoV-2 (wildtype) | ORF8:L84S |
| recombinant SARS-CoV-2 ΔORF6 | ORF8:L84S, ORF6: deletion (deleted positions: 27192-27385) |
| recombinant SARS-CoV-2 ORF6<sup>M58R</sup> | ORF8:L84S, ORF6:M58R (ATG to CGG) |
| BA.1 | GenBank: OL672836.1, Alias: B.1.1.529.1 |
| BA.2 | GenBank: OM371884.1, Alias: B.1.1.529.2 |
| BA.2.9.2 | GenBank: ON384332.1, Alias: B.1.1.529.2.9.2 |
| BA.4 | GenBank: ON373214.1, Alias: B.1.1.529.4 |
| BA.5 | GenBank: ON249995.1, Alias: B.1.1.529.5 |

**Table 2.**
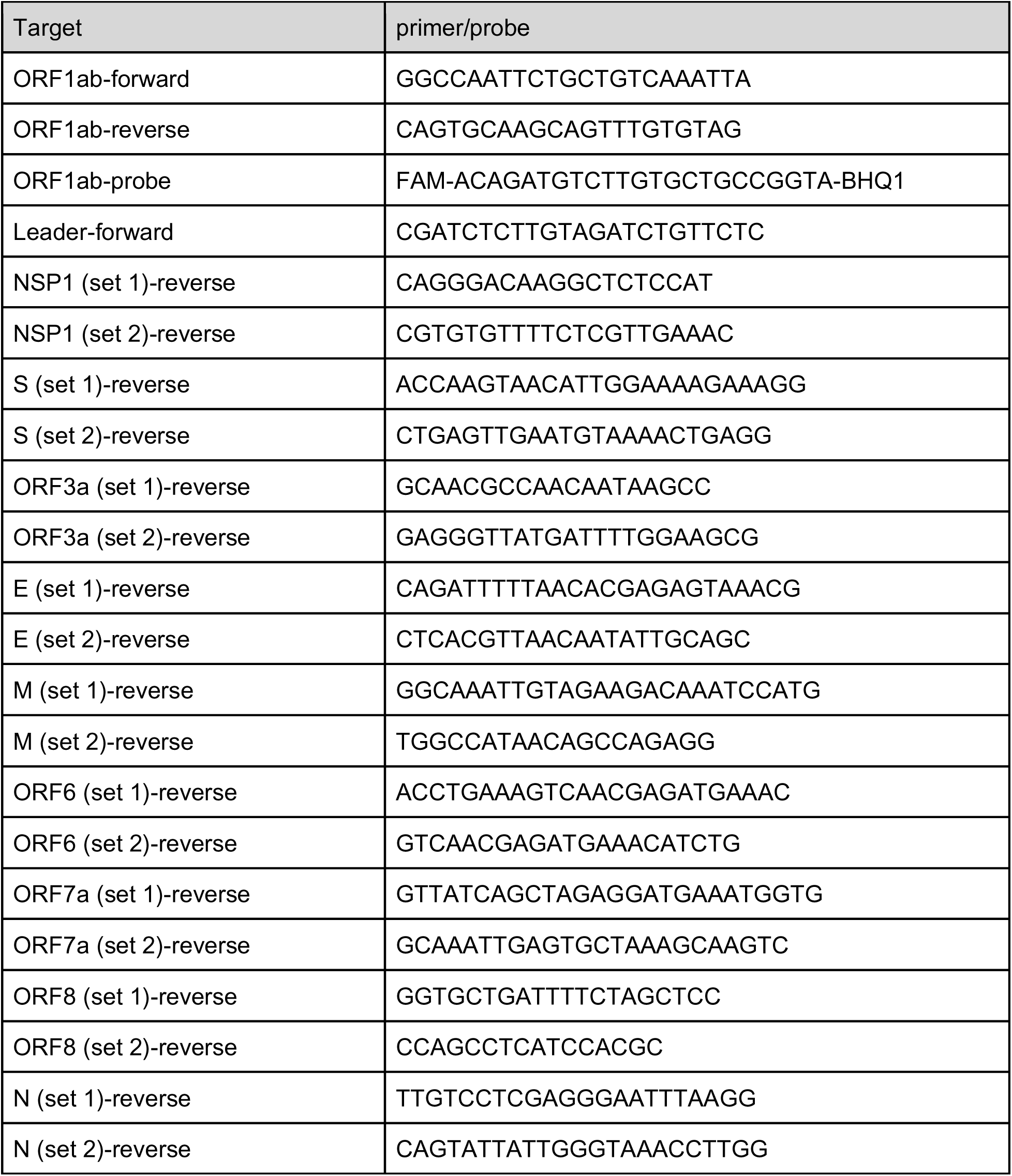
Primers and probes used in this study. Sequence given in 5’-3’ directionality.

### Immunolabeling with fluorescent in situ hybridization (Immuno-FISH)

HEK293T cells were seeded on glass-slides at a low density and transfected with the indicated plasmids for 24h. Cells were fixed, stained, and processed as described before (*11*). Confocal laser scanning microscopy was performed with a Zeiss LSM880 confocal laser scanning microscope (Carl Zeiss Microimaging) fitted with a Plan Apochromat 63×/1.4 or 40×/1.4 oil objective. Analysis of the nuclear/cytoplasmic ratio of Poly(A) RNA signal was performed as described elsewhere (*54*).

### Mass spectrometry (MS)

#### Cell lysis and digestion for proteomics

At the indicated time after infection A549-ACE2 cells were washed three times in ice cold 1x PBS. Next, cells were lysed in 500uL/well of 6M guanidine hydrochloride (Sigma) in 100mM Tris-HCl (pH 8.0) (Boston Bioproducts) and scraped with a cell spatula for complete collection of the sample. Samples were then boiled for 5 minutes at 95°C to inactivate proteases, phosphatases and the virus. Samples were frozen at −80°C until further processing. Samples were sonicated with a probe sonicator three times for 10 seconds at 20% amplitude. Insoluble material was pelleted by spinning samples at 13,000rpm for 10 minutes. Supernatant was transferred to a new protein lo-bind tube and protein was quantified using a Bradford assay. Samples were processed for reduction and alkylation using a 1:10 sample volume of tris-(2-carboxyethyl) (TCEP) (10mM final) and 2-chloroacetamide (4.4mM final) for 5 minutes at 45°C with shaking. Prior to protein digestion, the 6M guanidine hydrochloride was diluted 1:6 with 100mM Tris-HCl pH8 to increase the activity of trypsin and LysC proteolytic enzymes, which were subsequently added at a 1:75 (wt/wt) enzyme-substrate ratio and placed in a 37°C water bath for 16-20 hours. Following digestion, 10% trifluoroacetic acid (TFA) was added to each sample to a final pH ∼2. Samples were desalted under vacuum using 50mg Sep Pak tC18 cartridges (Waters). Each cartridge was activated with 1 mL 80% acetonitrile (ACN)/0.1% TFA, then equilibrated with 3 × 1 mL of 0.1% TFA. Following sample loading, cartridges were washed with 4 × 1 mL of 0.1% TFA, and samples were eluted with 2 × 0.4 mL 50% ACN/0.25% formic acid (FA). Approximately 60μg of each sample was kept for protein abundance measurements, and the remainder was used for phosphopeptide enrichment. Samples were dried by vacuum centrifugation. Thus, the same original sample was used for abundance proteomics and phosphoproteomics analysis.

#### Phospho-peptide enrichment for phospho-proteomics

IMAC beads (Ni-NTA from Qiagen) were prepared by washing 3x with HPLC water, incubating for 30 minutes with 50mM EDTA pH 8.0 to strip the Ni, washing 3x with HPLC water, incubating with 50mM FeCl3 dissolved in 10% TFA for 30 minutes at room temperature with shaking, washing 3x with and resuspending in 0.1% TFA in 80% acetonitrile. Peptides were enriched for phosphorylated peptides using a King Flisher Flex. For a detailed protocol, please contact the authors. Phosphorylated peptides were found to make up more than 90% of every sample, indicating high quality enrichment.

#### MS acquisition and data preprocessing for abundance proteomics

Digested samples were analyzed on an Orbitrap Fusion Lumos Tribrid mass spectrometry system (Thermo Fisher Scientific) equipped with an Easy nLC 1200 ultra-high pressure liquid chromatography system (Thermo Fisher Scientific) interfaced via a Nanospray Flex nanoelectrospray source. For all analyses, samples were injected on a C18 nano flow column (15 cm x 150 μm ID packed with PepSep1.9 μm particles). Mobile phase A consisted of 0.1% FA, and mobile phase B consisted of 0.1% FA/80% ACN. Peptides were separated by a linear gradient from 3% to 30% mobile phase B over 90 minutes, 30% to 38% B over 8 minutes, 38% to 88% B over 2 minutes, then held at 88% B for 10 minutes at a flow rate of 600 nL/minute (total of 110 minutes). Analytical columns were equilibrated with 6 μL of mobile phase A. Data was acquired using data independent acquisition (DIA) mode with the following parameters. A cycle consisted of a full FTMS scan at 120,000 resolving power over a scan range of 300-1400 m/z, a normalized AGC target of 100%, an RF lens setting of 30%, and a maximum injection time of 50 ms. DIA scan windows were variable, with 20 16m/z windows from 358-643m/z, 8 18m/z windows from 659-795m/z, 6 20m/z windows from 813-908m/z, 4 25m/z windows from 929.5-977.5m/z, 1 35m/z window at 1006.5m/z, 1 50m/z window at 1048m/z, and one 78m/z window at 1111m/z. Cycle time was 3 seconds. Loop control was set to 3. Raw mass spectrometry data from each run was analyzed the directDIA Analysis function in Spectronaut version 15.6.211220.50606 (Rubin) by Biognosys (no spectral library used). Data was searched against proteomics for Homo sapiens (downloaded February 28, 2020) and 29 SARS-CoV-2 protein sequences translated from genomic sequence downloaded from GISAID (accession EPI_ISL_406596, downloaded March 5, 2020). Data were searched using the default BGS settings, variable modification of methionine oxidation, static modification of carbamidomethyl cysteine, and filtering to a final 1% false discovery rate (FDR) at the peptide, peptide spectrum match (PSM), and protein level. Between run normalization was disabled and performed later using artMS (see below). On average, 5 data points per peak in MS1 and MS2 were captured per sample.

#### MS acquisition and data preprocessing for phosphoproteomics

Phospho-enriched samples were analyzed on a Q Exactive Plus Quadrupole-Orbitrap mass spectrometry system (Thermo Fisher Scientific) equipped with an Easy nLC 1200 ultra-high pressure liquid chromatography system (Thermo Fisher Scientific) interfaced via a Nanospray Flex nanoelectrospray source. For all analyses, samples were injected on a C18 reverse phase column (25 cm x 75 μm packed with ReprosilPur 1.9 μm particles). Mobile phase A consisted of 0.1% FA, and mobile phase B consisted of 0.1% FA/80% ACN. Peptides were separated by a linear gradient from 2% to 4% for 1 minute, 4% to 24% for 56 minutes, 24% to 38% for 19 minutes, 38% to 90% for 3 minutes, held at 90% for 8 minutes, then decreased from 90% to 2% for 1 minute and held at 2% for 2 minutes at a flow rate of 300nL/minute (total of 90 minutes). Analytical columns were equilibrated with 6 μL of mobile phase A. Data was acquired using data dependent acquisition (DDA) mode, acquired over a range of 300-1500 m/z in the Orbitrap at 70,000 resolving power with a normalized AGC target of 300%, an RF lens setting of 40%, and a maximum ion injection time of 60 ms. Dynamic exclusion was set to 60 seconds, with a 10 ppm exclusion width setting. Peptides with charge states 2-6 were selected for MS/MS interrogation using higher energy collisional dissociation (HCD), with 20 MS/MS scans per cycle. MS/MS scans were analyzed in the Orbitrap using isolation width of 1.3 m/z, normalized HCD collision energy of 30%, normalized AGC of 200% at a resolving power of 30,000 with a 54 ms maximum ion injection time. Raw mass spectrometry data from each run was analyzed using Maxquant (version 1.6.12). Data was searched against proteomics for Homo sapiens (downloaded February 28, 2020) and 29 SARS-CoV-2 protein sequences translated from genomic sequence downloaded from GISAID (accession EPI_ISL_406596, downloaded March 5, 2020). Data were searched using default settings, variable modification of methionine oxidation and phosphorylation (STY), static modification of carbamidomethyl cysteine, and filtering to a final 1% false discovery rate (FDR) at the peptide, peptide spectrum match (PSM), and protein level.

#### MS quantitative comparison analysis for abundance and phospho-proteomics

Quantitative analysis was performed in the R statistical programming language (version 4.0.2, 2020-06-22). Initial quality control analyses, including inter-run clustering, correlations, principal components analysis (PCA), peptide and protein counts and intensities were completed with the R package artMS (version 1.8.1). Based on obvious outliers in intensities, correlations, and clusterings in PCA analysis, 1 run was discarded from the protein abundance dataset (d6 [ΔORF6] replicate 2); no runs were discarded from the phosphorylation dataset. Statistical analysis of phosphorylation and protein abundance changes between wild-type (WT), d6 (ΔORF6), and M58R (ORF6M58R) infected samples were calculated using peptide ion fragment data output from Spectronaut and processed using artMS. Specifically, quantification of phosphorylation peptide ions were processed using artMS as a wrapper around MSstats, via functions artMS::doSiteConversion and artMS::artmsQuantification with default settings. All peptides containing the same set of phosphorylated sites (but different elution times or charge states) were grouped and quantified together into phosphorylation site groups. For both phosphopeptide and protein abundance MSstats pipelines, MSstats performs normalization by median equalization, imputation of missing values and median smoothing (Tukey’s Median Polish) to combine intensities for multiple peptide ions or fragments into a single intensity for their protein or phosphorylation site group, and statistical tests of differences in intensity between infected and control time points. When not explicitly indicated, we used defaults for MSstats for adjusted p-values, even in cases of N = 2. By default, MSstats uses Student’s t-test for p-value calculation and Benjamini-Hochberg method of FDR estimation to adjust p-values. After quality control data filtering, principal components analysis (PCA) and Pearson’s correlation confirmed strong correlation between biological replicates, time points, and conditions (except for the one run that was discarded).

#### Viral protein quantification

Median normalized peptide feature (peptides with unique charge states and elution times) intensities (on a linear scale) were refined to the subset that mapped to SARS-CoV-2 protein sequences as defined by MaxQuant (see above). Peptides found in the same biological replicate (i.e. due to different elution times, charge states, or modifications, for example) were averaged at the intensity level. Next, we selected the subset of peptides that were consistently detected in all biological replicates across all conditions (allowing no missing values), isolating the set of peptides that were consistently detected across all runs and thus possessing the best comparative potential. Isolating to this set of peptides, we summed all peptides mapping to each viral protein within each sample, which produced a final intensity value per viral protein, per sample. These resulting protein intensities were averaged across biological replicates and standard errors were calculated for each condition. To calculate the ratios, averaged intensities from each condition were divided (e.g. ΔORF6/WT). The standard error (SE) of the ratios was calculated as (A/B) * sqrt((se.A/A)² + (se.B/B)²).

### Bulk RNA Sequencing

Samples for bulk RNA sequencing were lysed in Trizol Reagent and total RNA was extracted using the miRNeasy mini kit (Qiagen) per the manufacturer’s instructions. DNAse treatment was performed on isolated RNA using the RNA Clean and Concentrator Kit (Zymo). Total RNA was examined for quantity and quality using the TapeStation (Agilent) and Quant-It RNA (ThermoFisher) systems. RNA samples with sufficient material (10 pg–10 ng) were passed to whole-transcriptome library preparation using the SMART-Seq v4 PLUS Kit (Takara Bio) following the manufacturer’s instructions. Briefly, total RNA inputs were normalized to 10ng in 10.5 µl going into preparation. 3’ ends of cDNA were then adenylated prior to ligation with adapters utilizing unique dual indices (96 UDIs) to barcode samples to allow for efficient pooling and high throughput sequencing. Libraries were enriched with PCR, with all samples undergoing 14 cycles of amplification prior to purification and pooling for sequencing. Bulk RNA sequencing was conducted on dual index libraries using a 300cycle Mid Output kit on an Illumina NextSeq 500 with standaer read configurations for R1, i7 index, i5 index, and R2:150, 8,8,150. Libraries were pooled and sequenced in two independent runs at 1.5 and 1.7pM loading concentrations. No PhiX was included in the loading library. Raw BCL files were converted to fastq files using bcl2fastq/2.20.0 (Illumina, Inc). For quantification of SARS-CoV-2 sgRNA and gRNA expression, the periscope/0.1.2 package was used with the technology argument set to “illumina” (*55*). Finally, sgRNA reads per total mapped reads were calculated. The SARS-CoV-2 reference was annotated with empirically identified SARS-CoV-2 sgRNAs (*56*).

### SARS-CoV-2 infection of Syrian Golden Hamsters

For the *in vivo* infection studies, experiments were conducted in 8-week-old female Syrian Golden Hamsters (Envigo, strain: HsdHan®:AURA) of approximately 120 grams body weight. The hamsters were housed in ventilated cages with free access to food and water and environmental enrichment. Cages were situated in a BSL3 vivarium with a light-cycle of 14 hours on, 10 hours off. Experimental protocols were approved by the Institutional Animal Care and Use Committee at Icahn School of Medicine at Mount Sinai (protocol number: IACUC-2017-0330). Hamsters were intranasally mock-infected (n=8) or infected with 5×10^5^ PFU of either rSARS-CoV-2 WT or rSARS-CoV-2 ΔORF6 (n=17 per group) in a 100uL total inoculum. Virus was diluted in PBS. Ketamine (100mg/kg)/ Xylazine(5mg/kg) was used to anesthetize the animals prior to infection. After infection, animals were monitored daily for morbidity and mortality up to day 15 post-infection. Necropsies were performed at 2, 4, 6, and 15 days post-infection (dpi). Animals were anesthetized with 200 uL Ketamine/Xylazine at a dose of 100 mg/kg of ketamine and 10 mg/kg xylazine and terminally bleed. Lungs and nasal turbinates were harvested. Total lung weight was measured. The left lobe of the lung was harvested, stored in Formalin (Fisherbrand), and processed for histology. The top right lobe of the lung was harvested and homogenized in Trizol. Total RNA was isolated with the RiboPure Kit (Invitrogen) per the manufacturer’s instructions. The bottom right lobe of the lung and nasal turbinates were homogenized in 750uL PBS and used for plaque assay. Matched hamsters were bled from the footpad at day 0 and day 15 post-infection and sera was isolated from whole blood by centrifugation to assess antibody titers. For the re-challenge experiment, animals were challenged with 1×10^5^ PFU of SARS-CoV-2 (USA-WA1/2020) (n=4 per group) at 30 days after initial infection. Animals were monitored for morbidity and mortality for up to 6 days post-challenge (36 days after initial infection). Nasal washes were performed at day 2, 4, and 6 post challenge with 250uL of PBS for assessment of viral titers by plaque assay.

### Immunohistochemistry (IHC)

#### Multiplex fluorescent immunohistochemistry

A Ventana Discovery Ultra (Roche, Basel, Switzerland) tissue autostainer was used for brightfield and multiplex fluorescent immunohistochemistry (fmIHC). In brief tyramide signaling amplification (TSA) was used in an iterative approach to covalently bind Opal fluorophores (Akoya Bioscience, Marlborough, MA) to tyrosine residues in tissue, with subsequent heat stripping of primary-secondary antibody complexes until all antibodies were developed. Lungs from infected (positive controls) and uninfected (negative controls) hamsters were used as controls for assay optimizing. In total two monoplex 3,3′-Diaminobenzidine (DAB) chromogenic assays (Ki67 and SARS-CoV-2 Spike) and two fluorescent duplexes: STAT1 + SARS-CoV-2 Spike, and MxA + SARS-CoV-2 Spike. Specific details for the immunohistochemical assays are outlined in Table 3, with a more concise overview provided below.

**Table 3.**
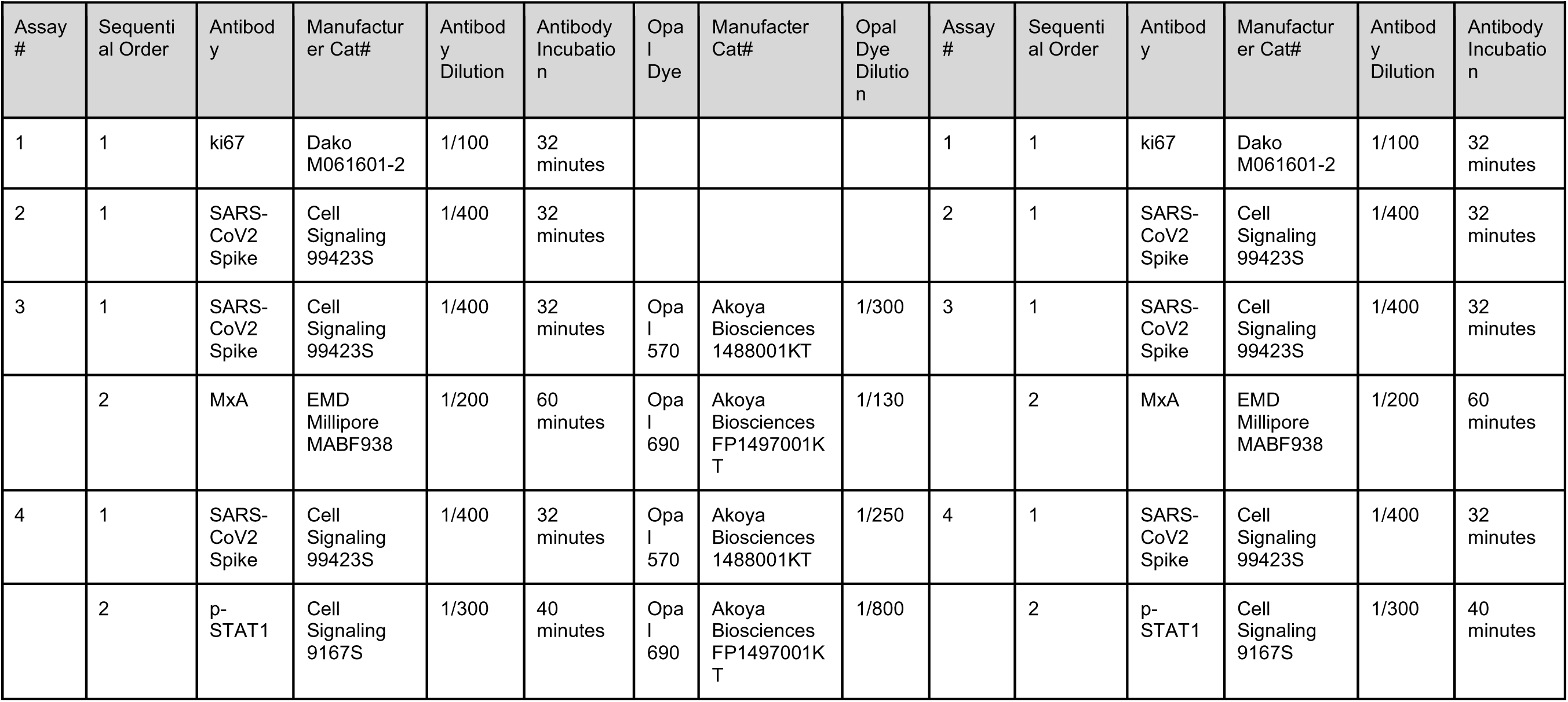
Supplemental information for IHC staining protocols.

#### Brightfield Immunohistochemistry

Antigen retrieval was conducted using a Tris based buffer-Cell Conditioning 1 (CC1)-Catalog # 950-124 (Roche). The SARS-CoV-2 spike primary antibody was of rabbit origin, and thus developed with a secondary goat anti-rabbit HRP-polymer antibody (Vector Laboratories, Burlingame, CA) for 20min at 37C. The Ki67 primary was of mouse origin, so a goat anti-mouse HRP-polymer antibody (Vector Laboratories) was utilized. Brightfield slides utilized A ChromoMap DAB (3,3′-Diaminobenzidine) Kit-Catalog #760-159 (Roche) to form a brown precipitate at the site of primary-secondary antibody complexes containing HRP. Slides were counterstained with hematoxylin and mounted.

#### Fluorescent Immunohistochemisty

Antigen retrieval was conducted using a Tris based buffer-CC1 (Roche). The SARS-CoV-2 Spike and Phospho-STAT1 primary antibodies were of rabbit origin, and thus developed with a secondary goat anti-rabbit HRP-polymer antibody (Vector Laboratories) for 20min at 37C. The MxA primary was of mouse origin, so a goat anti-mouse HRP-polymer antibody (Vector Laboratories) was utilized. All Opal TSA-conjugated fluorophore reactions took place for 20 minutes. Fluorescent slides were counterstained with spectral DAPI (Akoya Biosciences) for 16 minutes before being mounted with ProLong gold antifade (ThermoFischer).

#### Multispectral microscopy

Fluorescently labeled slides were imaged using a Vectra Polaris TM Quantitative Pathology Imaging System (Akoya Biosciences). Exposures for all Opal dyes on the Vectra were set based upon regions of interest with strong signal intensities to minimize exposure times and maximize the specificity of signal detected.

#### Digitalization and linear unmixing of multiplex fluorescent immunohistochemistry

Whole slide images were segmented into smaller QPTIFFs, uploaded into Inform software version 2.4.9 (Akoya Biosciences), unmixed using spectral libraries affiliated with each respective opal fluorophore including removal of autofluorescence, then fused together as a single whole slide image in HALO (Indica Labs, Inc., Corrales, NM).

#### Quantitative analysis of multiplex fluorescent immunohistochemistry

View settings were adjusted to allow for optimal visibility of immunomarkers and to reduce background signal by setting threshold gates on minimum signal intensities. Bronchioles, interstitium, and airways were classified using the tissue random forest tissue classifier module in HALO (Indica Labs), which was developed by annotating each tissue type via manual annotations. Separate layers for interstitium, bronchioles, and the whole lung were generated from the classifier, allowing algorithms to be ran on each layer for specific anatomical compartment analysis. These annotations were extensively examined for any errors by the machine-learning classifier and manually excised as necessary. For quantifying the area of the slide that contained SARS-CoV2 Spike, an algorithm called the HALO (Indica Labs) Area Quantification (AQ) module (v2.1.11) was created and finetuned to quantify the immunoreactivity for the Spike protein based on color and stain intensity. This algorithm outputted the % of total area displaying immunoreactivity across the annotated whole slide scan in micrometers squared (μm²). For quantifying the absolute number and overall percentage of cells expressing MxA we utilized the Halo (Indica Labs) HighPlex (HP) phenotyping modules (v4.0.4). In brief, this algorithm was used to first segment all cells within the annotated lung sections using DAPI counterstain. Detection threshold and nucleus geometry were defined until segmentation appeared accurate. Next, minimum nucleus, cytoplasm and membrane thresholds were set for each fluorophore to detect low and high expression within each of the segmented cells. Parameters were set using the real-time tuning mechanism that was tailored for each individual sample based on signal intensity. Phenotypes of infected MxA+, uninfected MxA+, infected MxA-, and uninfected MxA-cells were determined by selecting inclusion and exclusion parameters as follows respectively: MxA+S+, MxA+S-, MxA-S+, and MxA-S-. For quantifying the absolute number and overall percentage of Phospho-STAT1-expressing cells with SARS-CoV-2 infection, we utilized the Halo (Indica Labs) HighPlex phenotyping modules (v4.0.4). For determining cellular location of Phospho-STAT1 in infected cells, two algorithms were made. One captured the total number of infected cells expressing Phospho-STAT1 in the cytoplasm or nucleus, and the other determined the number of infected cells expressing STAT1 in the nucleus only (Fig. S6). HALO does not output specific cellular location counts of defined phenotypes, so two algorithms were necessary to determine cellular location within cells with more than one marker. By subtracting the number of nuclear-expressing Phospho-STAT1+ infected cells from the total Phospho-STAT1+ infected cells, the number of cytoplasmic-only expressing cells could be determined. Phenotypes of cells were determined by selecting inclusion and exclusion parameters as follows respectively: Spike+ Phospho-STAT1+, Spike+ Phospho-STAT1-, Spike-Phospho-STAT1+, and Spike-Phospho-STAT1-. By using the outputs of these two algorithms, the number of infected cells expressing Phospho-STAT1 in the cytoplasm only could be determined. The quantitative output for the AQ and HP was exported as a .CSV.

### Enzyme-Linked Immunosorbent Assay (ELISA)

96-well-microtiter plates (Thermo Fisher) were coated with 100 μL of recombinant spike protein of SARS-CoV-2 (Sino Biological, Cat. 40589-V08H4) at a concentration of 2 ug/mL at 4°C overnight. Plates were washed three times with PBS (Gibco) containing 0.1% Tween-20 (PBS-T) (Fisher Scientific) using an automatic plate washer (BioTek). After washing, plates were blocked for 1 hour at room temperature with 200 μL blocking solution per well (PBS-T with 3% milk powder (American Bio). After removing the blocking solution, serum samples were diluted to a starting concentration of 1:80, serially diluted 1:3 in PBS-T supplemented with 1% milk powder (American Bio) and incubated at room temperature for 2 h. The plates were washed three times with PBS-T and 100 uL anti-hamster IgG horseradish peroxidase antibody (HRP, abcam, #ab6892) diluted 1:10,000 in PBS-T containing 1% milk powder was added to all wells. After 1 hour of incubation at room temperature, plates were washed three times with 100 μL 3,3′,5,5′-Tetramethylbenzidine (TMB; Rockland, Cat# TMBM-100) using the plate washer and incubated at room temperature for 15 min. The reaction was stopped with 1 N sulfuric acid solution (Fisher Science). The absorbance was measured at 450 nm with a plate spectrophotometer (Synergy H1 hybrid multimode microplate reader, Biotek). Optical density (OD) for each well was calculated by subtracting the average background plus three standard deviations. Area under the curve (AUC) was computed using GraphPad Prism software.

---

**Supplementary Figure 1.**
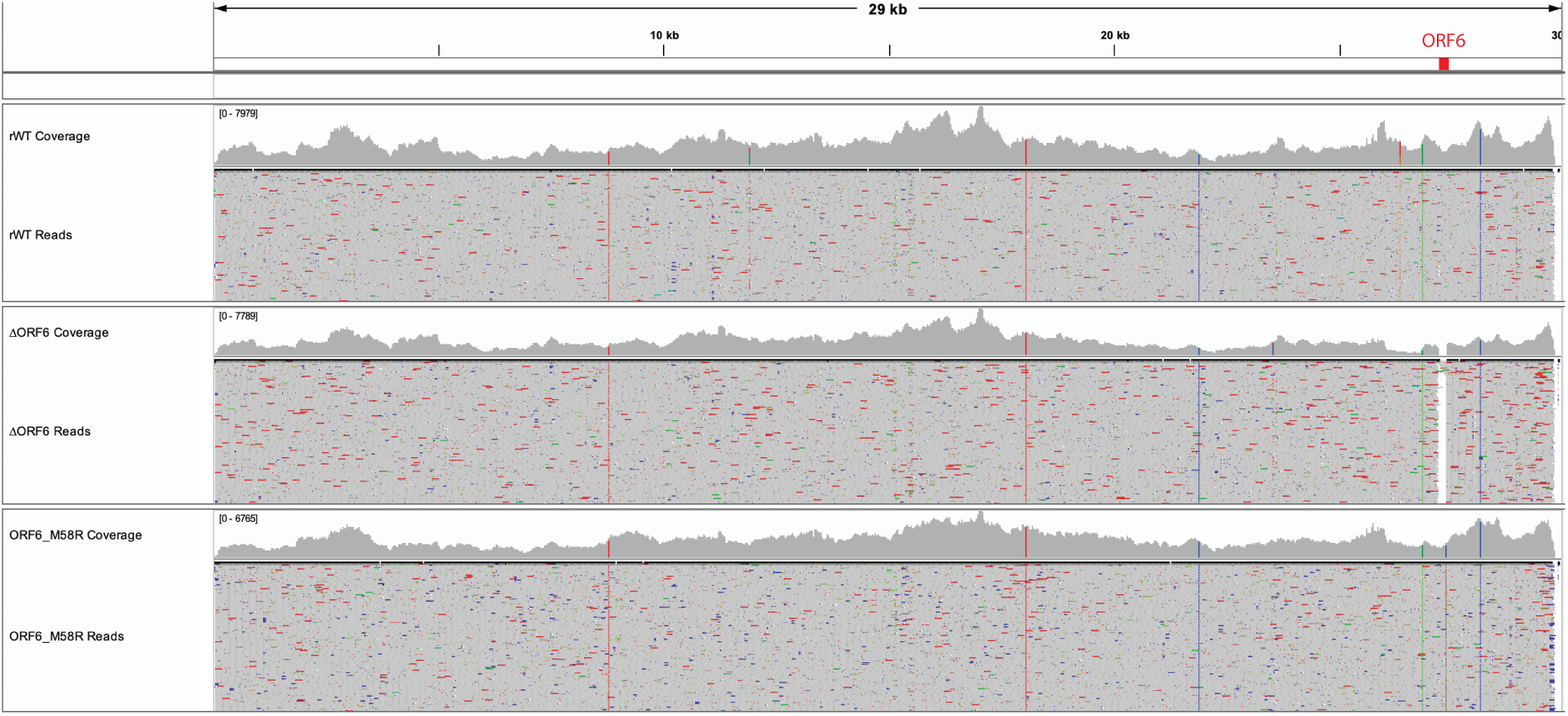
Sequencing of SARS-CoV-2 recombinant viruses. Deep sequencing data of RNA isolated from the indicated viral stocks confirming the presence of the expected deletion/mutations in ORF6. The graph shows aligned reads of all three viruses against a SARS-CoV-2 reference genome. The region encoding ORF6 is indicated in red. Grey indicates alignment to the reference, colorful lines indicate mutations.

**Supplementary Figure 2.**
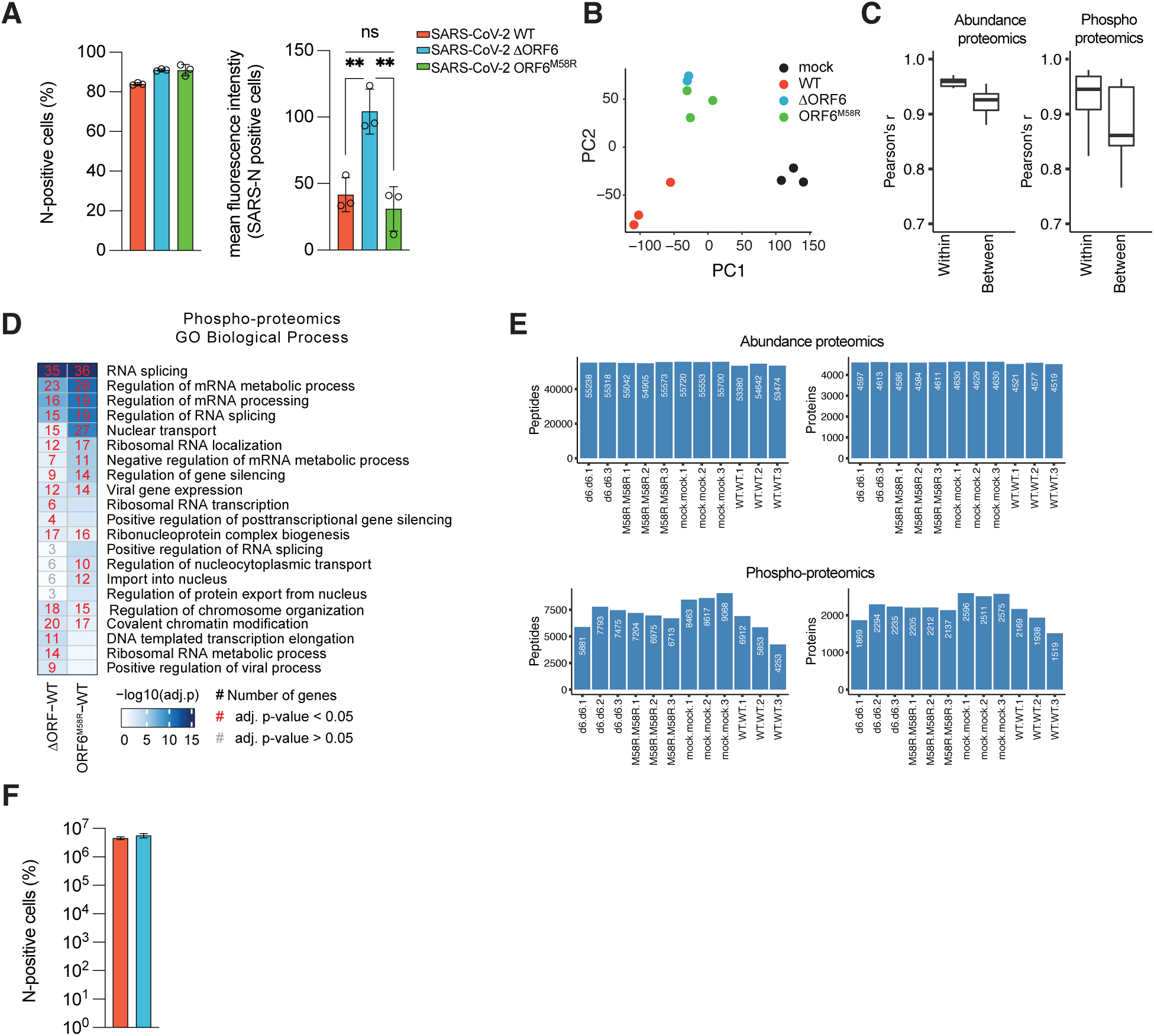
Proteomics quality control. (A) A549-ACE2 were infected with rSARS-CoV-2 WT, ΔORF6, or ORF6-M58R at MOI 2 for 24h before processing for flow cytometry as described in methods. Infection rates for each virus are shown by percentage of SARS-CoV-2 nucleocapsid positive cells, mean fluorescence intensity (MFI) is shown for N-positive cells (n=3). (B) Principal component analysis (PCA) of peptide intensities from abundance proteomics acquired during infection of cells and viruses described in A. (C) Pearson’s r correlation of peptide intensities from abundance proteomics (left) and phosphoproteomics (right) between biological replicates (“within” sample groups) and between conditions (“between” sample groups) for cells infected with viruses described in A. (D) GO Biological Process enrichment analysis of significantly differentially regulated (abs(log_2_FC)>1 & adjusted p<0.05) proteins from phosphoproteomics data obtained during infection with cells and viruses described in A. (E) Peptide (left) and protein (right) counts from each sample of the abundance proteomics (top) or phosphoproteomics (bottom) datasets obtained during infection of cells and virus described in A (F) A549-ACE2 were infected with rSARS-CoV-2 WT, ΔORF6, or ORF6-M58R at MOI 0.5 for 24h before processing for flow cytometry as described in methods. Infection rates for each virus are shown by percentage of SARS-CoV-2 nucleocapsid positive cells (n=3). Data in A and F were analyzed by ordinary one-way ANOVA using Turkey’s multiple comparison test. P > 0.05 = ns, P < 0.01 = **. Graphs were generated with PRISM (version 9).

**Supplementary Figure 3.**
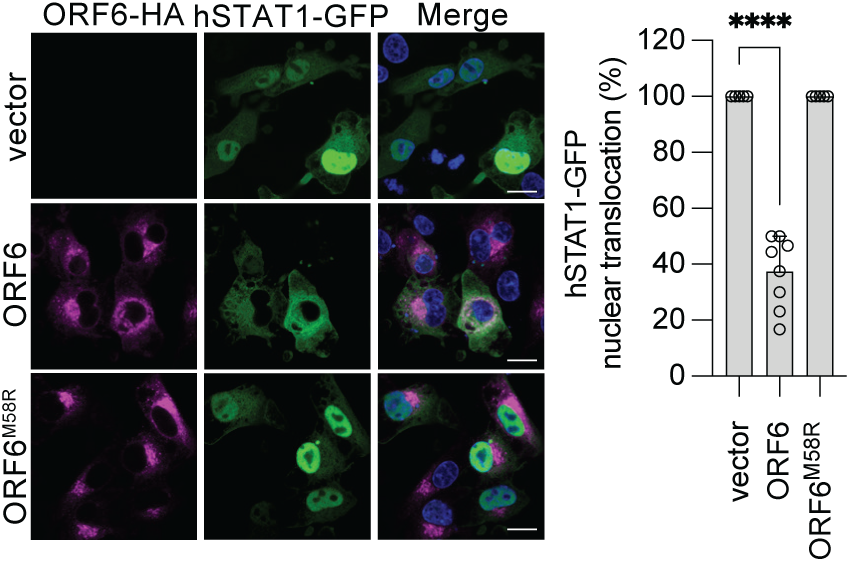
ORF6 antagonizes NUP98-Rae1-dependent STAT translocation in hamster cells. Confocal microscopy images of BHK-21 cells transfected with SARS-CoV-2 ORF6, ORF6-M58R or empty vector along with FLAG-RIG-I-2CARD and IRF3-GFP. At 24 h post-transfection, cells were fixed and processed for assessment of the subcellular localization of IRF3-GFP by confocal microscopy. Nuclear translocation of IRF3-GFP in control and ORF6/RIG-I-2CARD double-positive cells was quantified from three fields of view collected from two independent experiments. Data are shown as average ± SD. (Scale bar = 20 µm). Data were analyzed by ordinary one-way ANOVA using Turkey’s multiple comparison test. P < 0.0001 = ****. Graphs were generated with PRISM (version 9).

**Supplementary Figure 4.**
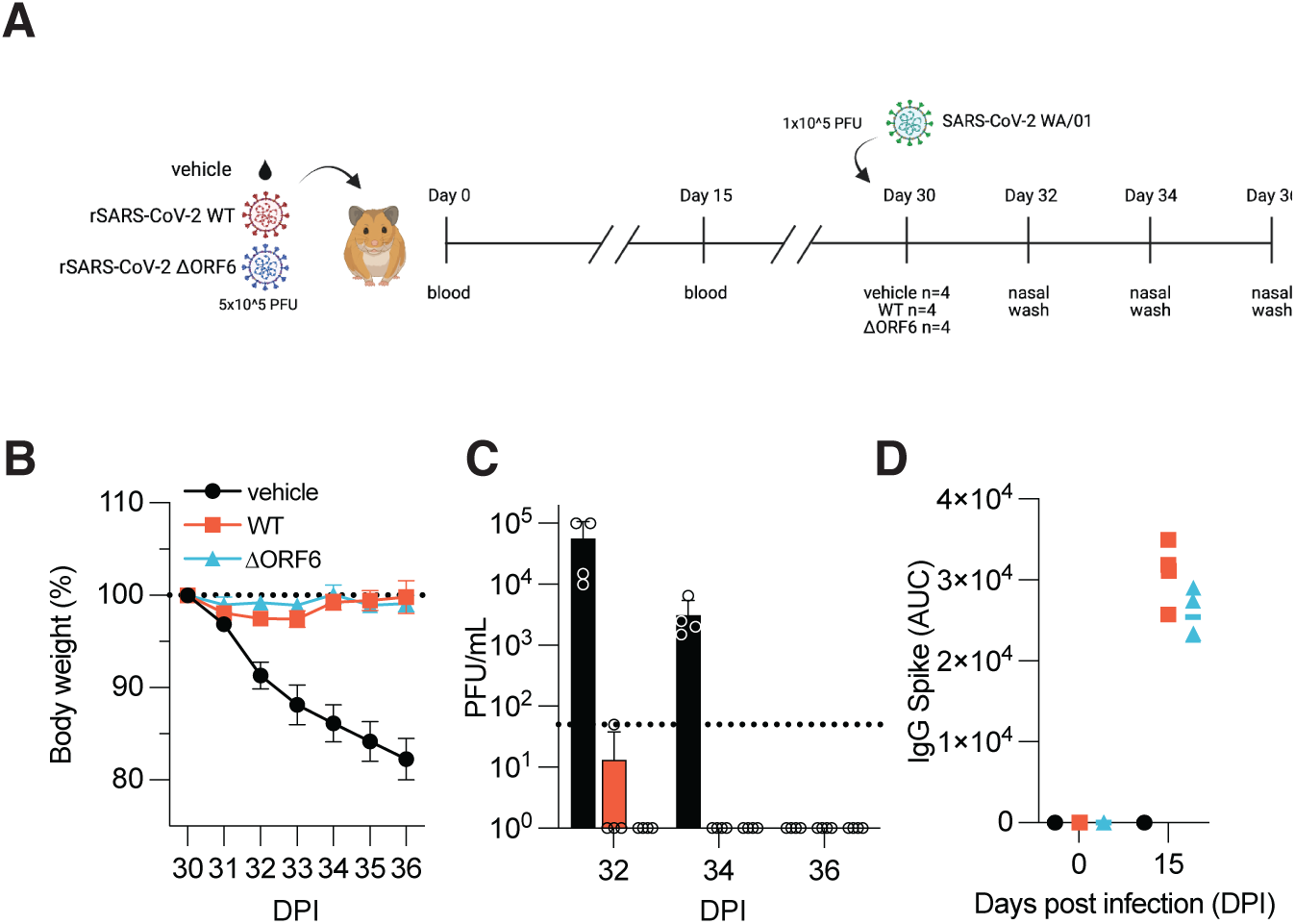
Infection with rSARS-CoV-2 ΔORF6 confers protection against challenge with the SARS-CoV-2 WA/01 isolate. (A) Schematic of the *in vivo* re-challenge experiment using recombinant SARS-CoV-2 viruses. In brief, hamsters previously infected with 5×10^5 PFU of the indicated viruses were subject to challenge with SARS-CoV-2 WA/01 30 days after initial infection and monitored for weight loss for 6 days post-challenge. Nasal washes were performed at 2, 4, and 6 days post-challenge to assess viral titers (n=4). (B) Weight loss curve for the duration of the challenge. Dashed line indicates 100 percent weight. Weight loss data is shown as mean ± SEM. (D) Nasal wash titers for animals at indicated days. Data is shown as PFU/mL. (C) Antibody titers of animals treated as described in A at 15 days after initial infection as measured by ELISA for Spike IgG. Data is showed as area under the curve (AUC). Data in D were analyzed by two-way ANOVA using Šídák’s multiple comparisons test.

**Supplementary Figure 5.**
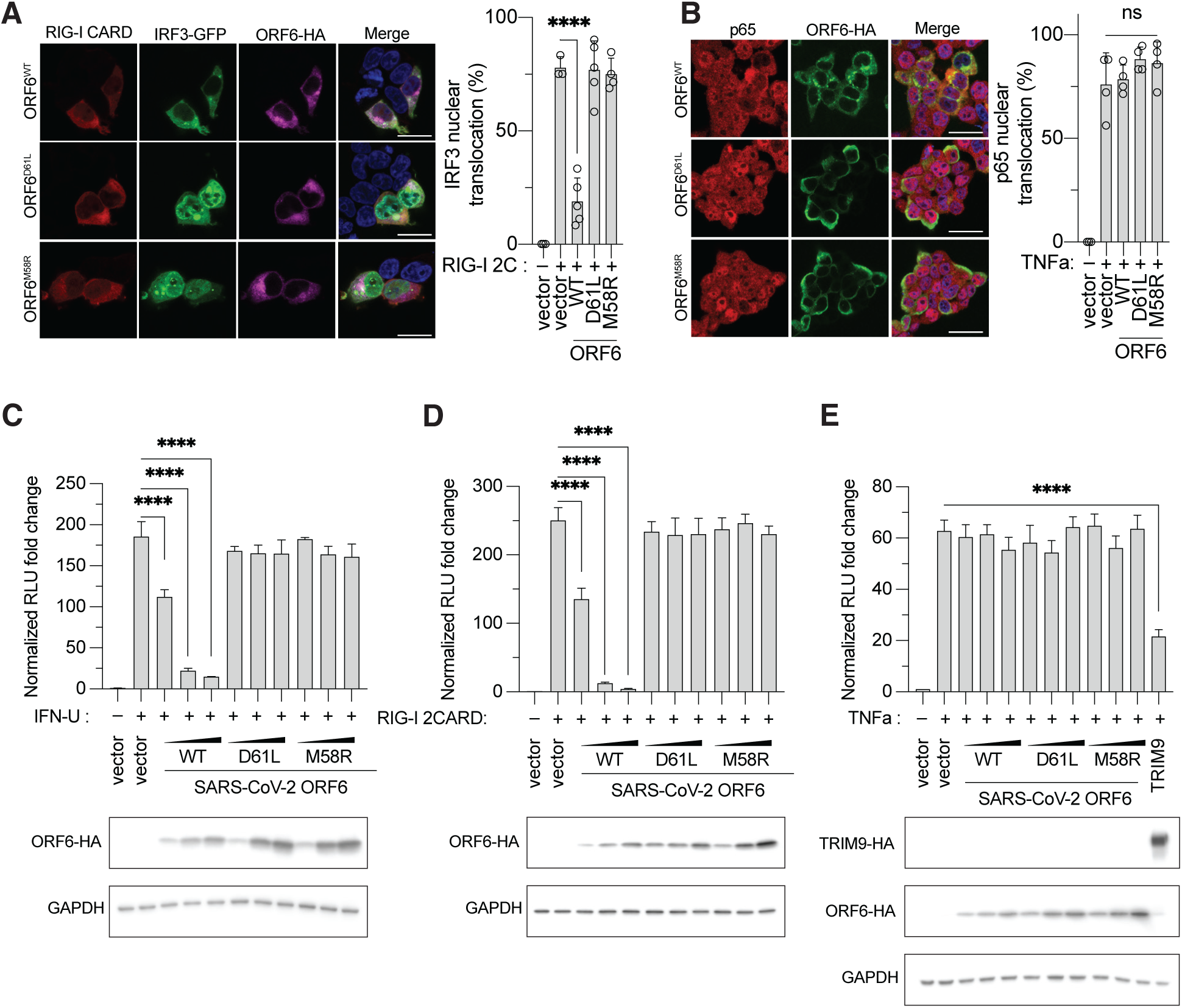
A D61L mutation in ORF6 disrupts protein functions. (A) Confocal microscopy images of HEK293T cells transfected with SARS-CoV-2 ORF6, ORF6-D61L, ORF6-M58R, or empty vector along with FLAG-RIG-I-2CARD and IRF3-GFP. At 24 h post-transfection, cells were fixed and processed for assessment of the subcellular localization of IRF3-GFP by confocal microscopy. Nuclear translocation of IRF3-GFP in control and ORF6/RIG-I-2CARD double-positive cells was quantified from four fields of view collected from two independent experiments. Data are shown as average ± SD. (B) Confocal microscopy images of HEK293T cells transfected with SARS-CoV-2 ORF6, ORF6-D61L, ORF6-M58R, or empty vector. At 24 h post-transfection, cells were treated with TNF-a (25ng/mL) for 45 min. Cells were fixed and processed for assessment of the subcellular localization of p65 by confocal microscopy. Nuclear translocation of p65 in control and ORF6-positive cells was quantified from four fields of view collected from two independent experiments. Data are shown as average ± SD. (C) HEK293T cells were transiently transfected with plasmids expressing ORF6, ORF6-D61L, or ORF6-M58R (0.5 ng, 2 ng, 5 ng, or 10 ng), a plasmid encoding an ISRE-firefly luciferase reporter, and plasmid expressing Renilla luciferase from the TK promoter. At 24h post-transfection, cells were treated with 1000 U of IFN universal for 16h, lysed and used for dual luciferase reporter assay. Data are representative of three independent experiments and shown as average ± SD (n = 3). Cell lysates from the reporter assay were analyzed by Western blot to show relative expression of each transfected viral protein. GAPDH was used as loading control. (D) HEK293T cells were transiently transfected with plasmids expressing ORF6, ORF6-D61L, ORF6-M58R (0.5 ng, 2 ng, 5 ng, or 10 ng), or HCV NS3/4A (50ng), FLAG-RIG-I-2CARD (5 ng), a plasmid encoding an 3xIRF3-firefly luciferase reporter (p55C1-luc), and plasmid expressing Renilla luciferase from the TK promoter. At 24h post-transfection, cells were lysed and used for dual luciferase reporter assay. Data are representative of three independent experiments and shown as average ± SD (n = 3). Cell lysates from the reporter assay were analyzed by Western blot to show relative expression of each transfected viral protein. GAPDH was used as loading control. (E) HEK293T cells were transiently transfected with plasmids expressing ORF6, ORF6-D61L, ORF6-M58R (0.5 ng, 2 ng, 5 ng, or 10 ng), or TRIM9 (100ng), a plasmid encoding an NFKB-firefly luciferase reporter, and plasmid expressing Renilla luciferase from the TK promoter. At 24h post-transfection, cells were treated with 25 ng/mL of TNF-a for 16h, lysed and used for dual luciferase reporter assay. Data are representative of three independent experiments and shown as average ± SD (n = 3). Lysates from the reporter assay were analyzed by Western blot to show relative expression of each transfected viral protein. GAPDH was used as loading control. (Scale bar = 20 µm). Data in A-D were analyzed by ordinary one-way ANOVA using Turkey’s multiple comparison test. P> 0.05 = ns, P < 0.0001 = ****. Graphs were generated with PRISM (version 9).

**Supplementary Figure 6.**
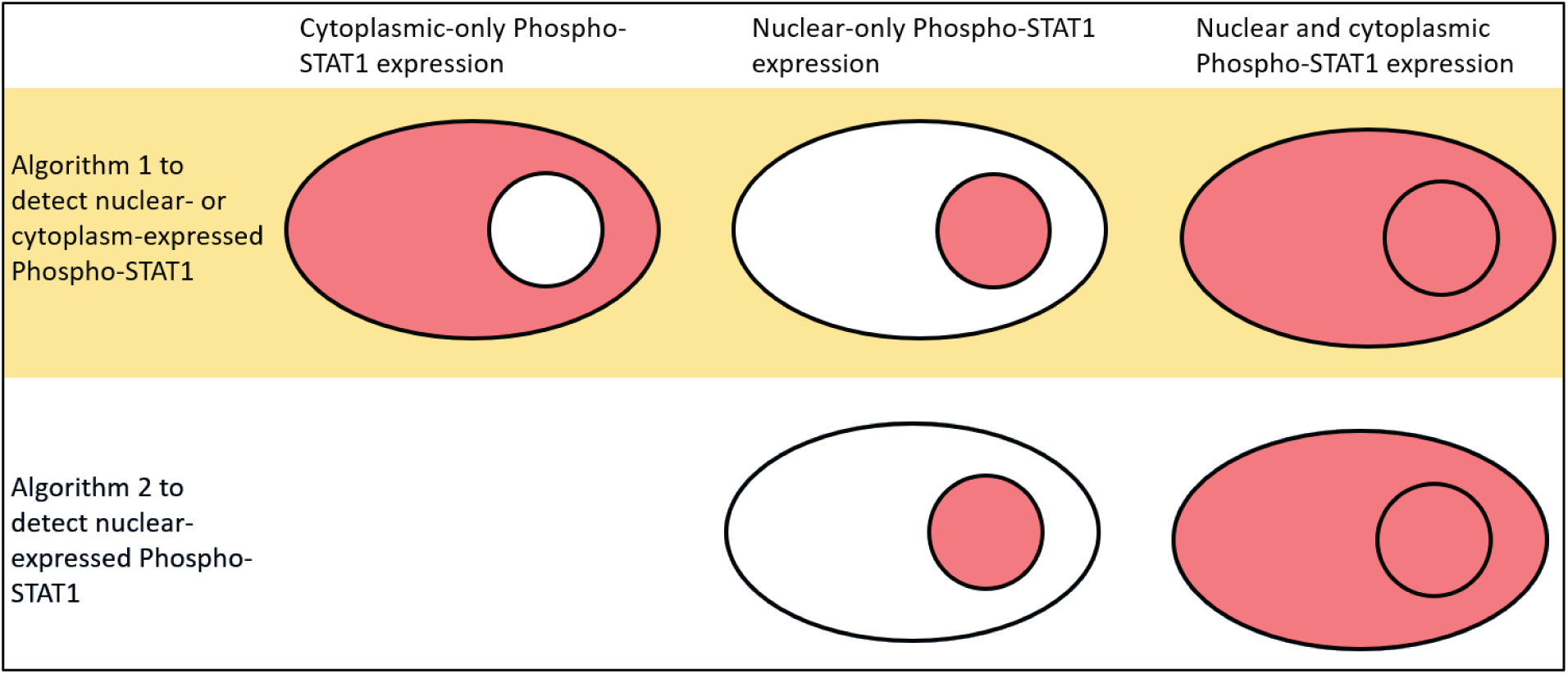
Schematic of algorithm for IHC quantification. Schematic of the algorithm created for quantification of nuclear pSTAT1 in SARS-CoV-infected cells in IHC stained lungs of infected Golden Syrian Hamsters.

